# The Hexosamine Biosynthetic Pathway alters the cytoskeleton to modulate cell proliferation and migration in metastatic prostate cancer

**DOI:** 10.1101/2024.10.14.618283

**Authors:** Rajina Shakya, Praveen Suraneni, Alexander Zaslavsky, Amit Rahi, Christine B. Magdongon, Raju Gajjela, Basil B. Mattamana, Dileep Varma

**Author notes:** Authors contributed equally (Rajina Shakya, Praveen Suraneni).

## Abstract

Castration-resistant prostate cancer (CRPC) progresses despite androgen deprivation therapy, as cancer cells adapt to grow without testosterone, becoming more aggressive and prone to metastasis. CRPC biology complicates the development of effective therapies, posing challenges for patient care. Recent gene-expression and metabolomics studies highlight the Hexosamine Biosynthetic Pathway (HBP) as a critical player, with key components like GNPNAT1 and UAP1 being downregulated in metastatic CRPC. GNPNAT1 knockdown has been shown to increase cell proliferation and metastasis in CRPC cell lines, though the mechanisms remain unclear.

To investigate the cellular basis of these CRPC phenotypes, we generated a CRISPR-Cas9 knockout model of GNPNAT1 in 22Rv1 CRPC cells, analyzing its impact on metabolomic, glycoproteomic, and transcriptomic profiles of cells. We hypothesize that HBP inhibition disrupts the cytoskeleton, altering mitotic progression and promoting uncontrolled growth. GNPNAT1 KO cells showed reduced levels of cytoskeletal filaments, such as actin and microtubules, leading to cell structure disorganization and chromosomal mis-segregation. GNPNAT1 inhibition also activated PI3K/AKT signaling, promoting proliferation, and impaired cell adhesion by mislocalizing EphB6, enhancing migration via the RhoA pathway and promoting epithelial-to-mesenchymal transition. These findings suggest that HBP plays a critical role in regulating CRPC cell behavior, and targeting this pathway could provide a novel therapeutic approach.

## Introduction

Prostate cancer is a type of cancer that begins in the cells of the prostate, and men over the age of 50, those with a family history of prostate cancer as well as African-American men are at a higher risk(1). Most prostate cancers initially respond to standard androgen deprivation therapy which is the standard treatment that reduces levels of male hormones, specifically testosterone(2). However, due to increased mutational load and genomic instability some tumors may develop resistance over time to this treatment, leading to castration-resistant prostate cancer (CRPC)(3). In CRPC, the cancer cells find ways to grow and survive. CRPC often indicates advanced disease state with a higher likelihood of metastasis. Despite low levels of circulating testosterone in CRPC, these cancer cells retain the ability to survive making treatment more challenging(4). Thus, a personalized and multidisciplinary approach, considering the unique characteristics of the nature of CRPC is crucial in managing complications and optimizing treatment outcomes.

The connection between metabolic pathways and cytoskeletal function plays a crucial role in tumor progression and metastasis in cancer(5). Cancer cells often undergo metabolic reprogramming to meet their increased energy and biosynthetic needs. A prominent example is the enhanced lipid metabolism, where increased fatty acid synthesis and oxidation support both rapid cell growth and metastasis(6,7). However, the mechanism by which metabolic pathways control prostate cancer is unclear(8). While it is known that this metabolic shift impacts the cytoskeleton through the regulation of actin dynamics, very little is known about how the microtubule cytoskeleton affect malignant properties, especially in prostate cancer(9). For example, elevated levels of fatty acid synthase (FASN) in prostate cancer have been linked to changes in cytoskeletal structure, which enhance cell motility and invasiveness(7). Additionally, prostate cancer cells often show increased glucose metabolism, which supplies key metabolites like ATP and acetyl-CoA to promote cytoskeletal reorganization. Rho GTPases, which regulate actin cytoskeleton remodeling, are frequently upregulated in prostate cancer, facilitating metastasis by promoting cell migration(5,7). These findings suggest that targeting metabolic pathways that influence the cytoskeleton may be a viable therapeutic strategy in treating advanced prostate cancer.

The hexosamine biosynthetic pathway (HBP) is a metabolic pathway that branches off from glycolysis and plays a crucial role in nutrient sensing and cellular regulation(10). The HBP is sensitive to nutrient availability and cellular stress, making it a crucial player in cellular responses to changing metabolic conditions(11). It involves the conversion of glucose to UDP-N-acetylglucosamine (UDP-GlcNAc), a key building block for glycosylation reactions. UDP-GlcNAc is involved in various cellular processes, including the modification of proteins through N/O-GlcNAcylation which regulates the function of many proteins, influencing processes such as signal transduction, transcription, and cell cycle progression. The HBP has been implicated in cancer, and its dysregulation is associated with various aspects of tumor development and progression(10–12). The HBP intersects with key cellular processes and signaling pathways, influencing cancer cells in several ways, including changes in glycosylation patterns, Insulin Resistance, and metabolic rewiring(13). Cancer cells often exhibit metabolic reprogramming, and dysregulated HBP contributes to altered cellular metabolism, providing advantages for tumor growth(8). The rate-limiting step of HBP involves the conversion of Gluocosamine-6-phosphate to N-acetylglucosamine-6-phosphate which is catalyzed by Glucosamine 6-phosphate N-acetyltransferase (GNPNAT1)(14). Recent studies suggest that the HBP pathway was downregulated in metastatic prostate cancer and knockout (KO) of GNPNAT1 induced cell proliferation and metastasis(8).

In this study, we hypothesize that the reduction in HBP components is responsible for alteration of actin and microtubule cytoskeleton as well as cell signaling pathways such as AKT and MAPK to control cell morphology, cell proliferation, and cell motility.

## Materials and Methods

### Cell lines and cell culture

The human normal prostate cell line RWPE1, human prostate cancer cell line DU145, and human CRPC cell line 22Rv1 were purchased from ATCC, and cultured in Dulbecco’s modified eagle’s medium (DMEM, Life Technologies) supplemented with 10% fetal bovine serum (Seradigm, VWR LifeScience), and 1% penicillin/streptomycin (Seradigm, VWR LifeScience). These wild-type cells along with the knockouts were grown at 37°C in a 5% CO_2_ humidified incubator.

### Chemical and reagents

Primary antibodies against GNPNAT1 and RND3 were purchased from ProteinTech whereas AKT, p-AKT, FOXO3a, p-FOXO3a, ERK, p-ERK, RhoA, RhoB, RhoC, PKCa, PKCzeta, PKCdelta, pPKCa/b, mTOR, p-mTOR, Raptor, Rictor, were purchased from Cell Signaling Technology (Denver, CO, USA). Antibodies against actin and tubulin were obtained from Santa Cruz Biotechnology (Dallas, TX, USA). Antibodies against MAP10 was purchased from LSBio (Lynnwood, WA, USA), and GAPDH from Millipore (Burlington, MA, USA). EphB6, ECAD, and NCAD from Abclonal (Woburn, MA, USA). MK2206 was purchased from AdooQ Bioscience (Irvine, CA) and UDP-GlcNAc from Millipore sigma (Burlington, MA).

### CRISPR/Cas9-mediated Knockout

The gRNAs targeting the human GNPNAT1 gene in 22Rv1 cells were designed using CRISPR based Gene Knockout kit v2 (Synthego Corporation). The sequences of different multiguide RNAs were: U*A*U*UUGAACAAAAACAGAAG,U*A*C*UUACUCAUAAAUUGUUC,U*U*C*AAAACUAGGUUUU UUUA. 22Rv1 CRPC cells were seeded and propagated to desired confluency in a 37°C incubator with 5% CO_2_. The GNPNAT1 gRNAs were incubated with spCas9 2NLS protein for 20 min at RT to make the complex of Cas9 RNPs. The RNP complex was then electroporated into 22Rv1 cells using the Neon NxT Electroporation System (ThermoFisher Inc.) under the kit conditions provided by the manufacturer, Synthego. For positive control, the same reaction was performed with a Synthego control gRNA, which was not specific to the human genome. The electroporated cells were cultured for 24-48 h and sorted as singles cells into 96 well plates containing DMEM medium supplemented with 20% FBS. We then monitored the cell growth of single colonies for 4-6 weeks while the cells grew into colonies, followed by scaling up into 24 well, and then into 6 well plates. The cells were then split into two groups, one for cryopreservation and the other to validate by western blot to confirm homozygous knockout clones.

### Immunofluorescence microscopy

Cells were fixed onto coverslips either with 100% ice-cold methanol or 3.6% paraformaldehyde. Coverslips were first rinsed three times in 1x PBS. For methanol fixation, cells were pre-fixed with ice-cold methanol for 1 min, then incubated at −20℃ for 5-6 mins. Alternatively, cells were permeabilized with 0.5% Triton X-100 for 5 mins, then fixed with 3.6% paraformaldehyde for 20 mins at room temperature (RT). Blocking was performed with 0.1% BSA in PBS for 1 h at RT. Cells were immunostained with primary antibody overnight at 4℃, washed three times with 1x PBS for 10 mins each, then incubated with Alexa Fluor-488/647 or Rhodamine Red-X (Thermo Fisher Scientific) secondary antibody, as required, for 1 h at 37℃. After two more washes in PBS, nuclei/chromosomes were counterstained with DAPI in 1x PBS (1:10,000) for 10 mins at RT and the coverslips were mounted onto glass slides with ProLong Gold Antifade reagent (Thermo Fisher Scientific) mounting media. Slides were stored at 4℃ until visualization.

### Image acquisition and processing

For fixed-cell imaging, three-dimensional stacks were captured using a Nikon Eclipse TiE inverted microscope, equipped with a Yokogawa CSU-X1 spinning disc, an Andor iXon Ultra888 EMCCD camera, and either a x60 or x100 1.4 NA Plan-Apochromatic DIC oil immersion objective (Nikon). During fixed-cell experiments, images were acquired at room temperature (RT) as Z-stacks with 0.2-0.3 µm intervals, controlled by NIS-Elements software (Nikon). The images were processed in NIS-Elements and presented as maximum-intensity projections of the required z-stacks. The intensities of the acquired images were quantified through ImageJ Fiji, and the corresponding data were plotted into graphs using Origin 2018 software for further analysis.

### Cell spreading assay

Live-cell imaging of cell spreading assay in control and GNPNAT1 KO 22Rv1 cells was carried out on 35-mm glass-bottomed dishes (MatTek Corporation) in an incubation chamber for microscopes (Tokai Hit Co., Ltd) at 37°C and 5% CO_2_ using FluoroBrite DMEM live imaging media (Life Technologies) supplemented with 2.5% FBS. However, in this case, the 35-mm glass-bottom dishes were manually coated with 10 µg/ml fibronectin prior to using them for cell culture. Images were acquired every 15min for up to 6 h, as required based on whether the samples were control or GNPNAT1 KO 22Rv1 cells.

### Western Blotting

The cells were harvested, washed with PBS, and lysed with RIPA buffer (Sigma-Aldrich, #R0278) containing Halt Protease Inhibitor Cocktail (Thermo Scientific, #87786) and incubated on ice for 20 min. Samples were centrifuged at 14,000 rpm at 4℃ for 5 min and the supernatant was collected. Protein concentrations were determined with Coomassie protein assay, then samples were prepared with Laemmli Sample Buffer and boiled at 100℃ for 10 min. Samples were resolved on 12-15% SDS-PAGE gel, then transferred to PVDF blotting membrane (Cytiva Amersham Hybond, #10600023).

For western blotting, PVDF membrane was blocked in 5% non-fat dry milk made in 1x TBS with 0.1% Tween20 solution (TBST) for 1 h at RT with shaking, followed by three 10 min washes with 1x TBST. Primary and secondary antibodies were prepared in 1x TBST with 5% BSA. The membranes were incubated at RT shaking with appropriate primary antibodies for 1 h, washed three times with 1x TBST, then incubated with goat anti-rabbit (Azure Biosystems, #AC2114) or goat anti-mouse HRP secondary antibodies (Azure Biosystems, #AC2115) used at 1:2000 for 1 h shaking at RT. Detection was achieved using SuperSignal West Pico PLUS Chemiluminescent Substrate (Thermo Fisher Scientific, #34580).

### Cell proliferation assay

The cell proliferation of 22Rv1, RWPE1, and DU145 cells was assessed using the Cell Proliferation Reagent WST-1 kit. Briefly, 10⁴ cells were seeded into quadruplicate wells of a 96-well plate and incubated at 37 °C under 5% CO₂/95% air for the specified period. WST-1 reagent was prepared as a five-fold dilution in the media. Then, 200 µL of WST-1 solution were added to each well, and the plate was incubated for an additional 2 hours. Subsequently, the optical density of the wells was measured at 450 nm using a Promega microplate reader.

### RNA sequencing analysis

RNA was prepared from WT and GNPNAT1 KO using the Direct-zolTM RNA MiniPrep kit from Zymo Research (#R2050). RNA quality control was performed at the NUSeq Core facility at Northwestern University. Library preparation, sequencing, and data analysis (Standard Analysis Package) was performed using the Standard RNA-seq services from GENEWIZ, Inc. The Volcano plot depiction of the analyzed data was generated using the R Program.

### Metabolomics analysis

Control and GNPNAT1 KO cell samples were prepared and send to Beth Israel Deaconess Medical Center-Mass spectrometry core facility for metabolomic analysis as described previously in Yuan et al. Briefly; to extract metabolites from control and GNPNAT1 KO cell lines, medium was replaced 2 hours before extraction. Cells are then washed and placed on dry ice, followed by the addition of 4 ml of 80% methanol at −80 °C. After incubating for 20 minutes at −80 °C, cells are scraped and transferred to conical tubes on dry ice. The mixture is centrifuged at 14,000g for 5 minutes at 4–8 °C to separate the debris, and the supernatant containing metabolites is collected. An additional methanol wash is performed on the pellet, followed by another centrifugation, and the supernatant is pooled. The total extract is divided into aliquots and dried using a SpeedVac. The metabolite extracts are then prepared for LC-MS/MS analysis by resuspending in LC/MS-grade water and injecting 5–10 µl onto the system. Data are acquired in SRM mode, and peaks are integrated using software like MultiQuant to quantify metabolite levels(15).

### Glyco-proteomics analysis

Immunoprecipitation of N-glycosylated proteins: To a 1000 μl Cerebellar lysate (1 μg/μl), 20 μg/ml biotinylated Concanavalin (Con-A) was added, and the reaction mixture was incubated overnight at 4°C rotating end over end. NeutrAvidin agarose beads (29204, ThermoFisher, Scientific-25 μl) were used to recover Con-A bound proteins at 4°C for 1 h. The mixture was spun at 2500 g for 2 min to collect the beads. The beads were washed with PBS 3 times followed by elution of N-glycosylated proteins in 2.5X SDS buffer by boiling the samples at 95°C for 10 min. After immunoprecipitation, N-glycosylated proteins were sent to Northwestern University Proteomics Core facility where Samples were precipitated overnight using eight volumes of acetone and one volume of TCA. The resulting protein pellet was resuspended and vortexed in 100 µL of 8 M urea and 0.4 M ammonium hydrogen bicarbonate (AmBic) solution. Reduction was performed by adding 4 µL of 100 mM dithiothreitol (DTT) and incubating for 30 minutes at 55 °C. Alkylation was carried out in the dark using 18 mM iodoacetamide for 30 minutes at room temperature. The urea concentration was then reduced to 1.8 M by adding four volumes of water. Digestion occurred overnight at 37 °C using MS-grade trypsin (Promega, Madison, WI) at an enzyme-to-substrate ratio of 1:50. To halt the digestion, 10% formic acid was added to achieve a final concentration of 0.5%. The peptides were desalted using Pierce C-18 spin columns and eluted with 80% ACN/0.1% FA, then dried in a vacuum concentrator. The dried peptides were resuspended in 30 µL of 5% ACN/0.1% FA for LC-MS analysis with a Dionex UltiMate 3000 Rapid Separation nano-LC paired with a Q Exactive HF (QE) Quadrupole Orbitrap mass spectrometer (Thermo Fisher Scientific Inc., San Jose, CA, USA). Identified peptides and proteins were visualized using Scaffold software (version 5.0, Proteome Software Inc., Portland, OR). The data were searched against a particular database using the MaxQuant application. For statistical analysis, a t-test was performed with a significance threshold of p < 0.05 and a fold change (FC) greater than 2 to identify significantly up- and down-regulated proteins, which were visualized using a volcano plot.

### Nuclear and cytosolic fractionation

The isolation of nuclear and cytosolic fractions of control and GNPNAT1 KO 22Rv1 cells was carried out based on the protocol(16). Initially, cells were harvested and washed with cold PBS. Cells were then resuspended in STM buffer comprising 250 mM sucrose, 50 mM Tris-HCl pH 7.4, 5 mM MgCl_2_, protease, and phosphates inhibitor cocktails, and homogenized for 1 min on ice using a Teflon pestle in Potter S homogenizer and vortexed. Following that, cells were centrifuged at 800 g for 15 mins. The supernatant was used for isolation of cytosolic fraction and the pellet was used for nuclear fraction. The pellet was further washed in STM buffer, vortexed at maximum speed and centrifuge at 500 g for 15 min. Supernatant was discarded and the washed pellet was resuspended in NET buffer comprising 20 mM HEPES pH 7.9, 1.5 mM MgCl_2_, 0.5 M NaCl, 0.2 mM EDTA, 20% glycerol, 1% Triton-X-100, protease, and phosphatase inhibitors.

For cytosolic fraction, above mentioned supernatant was further centrifuged at 11,000 g for 10 min and the supernatant was precipitated in cold 100% acetone at -20°C for at least 1 h followed by centrifugation at 12,000 g for 5 mins. The pellet containing cytosolic fraction was then resuspended in STM buffer.

### Proteome array analysis

Proteome array analysis was conducted for Human Phospho-Kinase proteome array according to the manufacturer’s instruction. In brief, cells were harvested and lysed with ice cold lysis buffer 17 provided with the Human Phospho-RTK proteome array kit (Bio-techne, Minneapolis, USA) for 30 min on ice. The samples were then centrifuged at 14,000 rpm for 10 min, supernatant was collected, and quantification of protein was carried out using Coomassie protein assay. The arrays were then blocked for 1 h by Array buffer 1 included in the kit, which serves as a blocking buffer. Then, they were incubated with 600 μg protein samples overnight at 4°C. The unbounded proteins on array were removed by washing three times with washing buffer. Arrays were incubated with anti-phospho-tyrosine-HRP detection antibody for 2 h at RT on a rocking platform shaker followed by three times washing with wash buffer. After the final wash, the excess wash buffer was allowed to drain from the array and Chemi Reagent Mix was added evenly onto each membrane. Multiple exposures were carried out to visualize the protein spot and the average intensity was calculated using ImageJ software program (Bethesda, Maryland, USA). After subtracting the averaged background signal, the fold change was obtained by comparing GNPNAT1 KO cells with control 22Rv1 cells.

### Wound healing assay

The wound healing assay was conducted using the Ibidi culture-insert 2-well system (Lochhamer Schlag, Germany). Initially, cells were harvested and seeded at a final concentration of 3×10⁵ cells/ml, then incubated at 37°C with 5% CO₂ for 24 hours. After incubation, the culture insert was carefully removed, and pre-warmed, cell-free medium (37°C) was gently added by pipetting. The cells were then allowed to migrate, and their movement was monitored using phase contrast microscopy. Imaging began with the acquisition of single frames at various time points (0, 6, 24, 48, and 72 hours) depending on the rate of migration.

### Cell migration and invasion assay

Initially, the inner surface of the Transwell insert (BD Falcon, Franklin Lakes, USA) was coated with collagen (1 mg/ml) for the cell migration assay and with Matrigel (0.5 mg/ml) for the cell invasion assay. Following coating, 100 μl of cell suspension (5×10⁵ cells/ml) in serum-free media was added to the inner chamber of the Transwell and incubated at 37°C with 5% CO₂ for 48 hours. After the 48-hour incubation, migrated and invaded cells were treated with the WST cell proliferation assay kit, and absorbance was measured at 440 nm using a Promega microplate reader.

### Statistical analysis

Data are presented as the mean standard deviation. Single comparisons were performed using the student’s t-test. Graphs and data plots were designed using Origin 2018 software. A probability (p) value less than 0.05 was considered statistically significant.

## Results

### HBP depletion affects various genes, metabolites, and glycoproteins altering the associated components and pathways

Previous studies employing cellular phenotypic analysis, and a network-based integrative approach have shown that specific alterations in the HBP are crucial for metastatic cancers such as CRPC. As published work has demonstrated, we consistently find a > 60% reduction in GNPNAT1 in 22RV1 CRPC cells as compared to RWPE1, a normal prostate cell line (Figure 1A). According to this study, the HBP pathway was downregulated in CRPC, which may contribute to the increased cell proliferation and metastasis(8). However, the physiological basis of these cancer-associated phenotypes is unclear. In order to achieve a more thorough and comprehensive understanding of the function of the HBP in cellular properties, we generated a CRISPR-mediated GNPNAT1 KO in 22Rv1 CRPC cell line.

**Figure 1.**
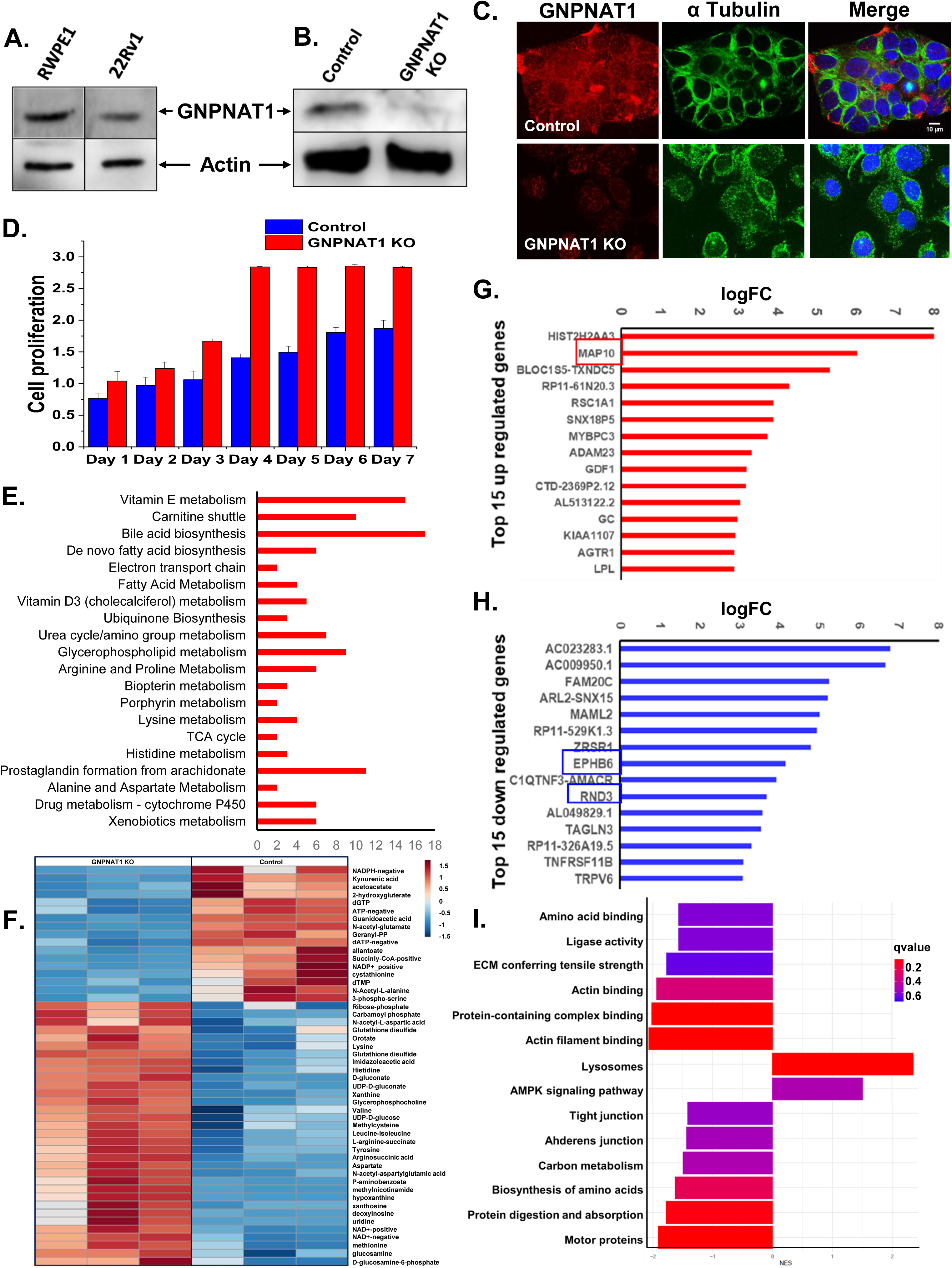
HBP depletion alters various genes, metabolites, glycoproteins as well as their associated pathways and processes. A) Western blot showing the downregulation of GNPNAT1 in CRPC (22Rv1) cells as compared to normal (RWPE1) cells. (B, C) Confirmation of CRISPR CAS9-mediated knockout of GNPNAT1 in 22Rv1 cells by (B) western blotting and (C) immunofluorescence staining. Scale bar, 10 µm. (D) Cell proliferation assay of control and GNPNAT1 KO 22Rv1 cells at days 1-7. (E) Depiction of the top 20 metabolic pathways that were significantly altered after GNPNAT1 inhibition. (F) Heat map representation of the altered metabolites after GNPNAT1 inhibition using metabolites expression analysis of control and GNPNAT1 depleted 22Rv1 cells. (G, H) Representation of the top 15 down-regulated (G) and up-regulated genes (H) via differential gene expression analysis of GNPNAT1 depleted 22Rv1 cells. (I) Total proteomic analysis of control and GNPNAT1 depleted 22Rv1 cells showing the pathways and processes altered.

We show the successful knockout of GNPNAT1 by our CRISPR-mediated approach, which was confirmed by both western blotting and immunofluorescence (Figure 1B and 1C). As shown by the published work, we found that starting from day 3, cells start to exhibit increased cell proliferation in GNPNAT1 KO cells as compared to the control 22Rv1 cells (Figure 1D). Using the same approach, we knocked out GNPNAT1 in RWPE1, a normal human prostate cell line and DU145, a human prostate cancer cell line (Supplementary Figure S1A), where we also observed the increased cell proliferation in the GNPNAT1 KO cells from day 3 (Supplementary Figure S1B-C).

In order to investigate and better comprehend the cellular phenotype in GNPNAT1-inhibited cells, we carried out transcriptomics analysis in cells depleted of GNPNAT1, since there is limited data available concerning how GNPNAT1 inhibition affects various cellular phenotypes. Our initial transcriptomic analysis allowed for a deeper understanding of the molecular mechanism driving cell proliferation, offering insights into normal growth and potential abnormalities associated with HBP-inhibited castration-resistant prostate cancers. This transcriptomic analysis along with gene ontology analysis (Supplementary Figure S2A-E) revealed the involvement of HBP with the alteration of expression of several genes, cellular components, and biological processes.

We used differential gene expression analysis in order to identify the genes that are specifically associated with the depletion of GNPNAT1. About 994 genes were found to be differentially expressed in the transcriptome of GNPNAT1 knockdown (KD) cells, indicating a major impact on gene expression in general (Supplementary Figure S2A). Several cell signaling genes and genes linked to cytoskeletal components were found to be enriched, according to this analysis (Figure 1G and 1H). Among these genes, we were interested in the significant upregulation of the microtubule-binding protein gene MAP10 and the downregulation of the genes EPHB6 and RND3, which will be addressed in more detail in this study.

Gene Ontology Analysis of biological processes suggest that several gene groups closely associated with cell adhesion and migration were severely dysregulated (Supplementary Figure S2B-E). GNPNAT1 KD 22Rv1 cells showed markedly altered pathways, including upregulated cell migration and downregulated extracellular matrix organization and disassembly, cell-cell adhesion, and glycosaminoglycan metabolism (GAGs). Based on this finding, we hypothesized that modifications to the HBP pathway could possibly impact the cell adhesion and migratory properties of CRPC cells. Numerous cytoskeletal components were also found to be dysregulated when we performed gene ontology analysis of cellular components (Supplementary Figure S2B-C). In accordance with that, we noticed that the GNPNAT1 KO cells had dramatically altered actin and microtubule cytoskeletons. When we compared the immunostaining and their intensity for GNPNAT1 KO 22Rv1 cells to the control, we observed that both tubulin and actin were significantly downregulated. In the control, tubulin and actin were present throughout the cytosol, however in the GNPNAT1 KO 22Rv1 cells, they were only present towards the cell periphery (Figure 2A-C).

**Figure 2.**
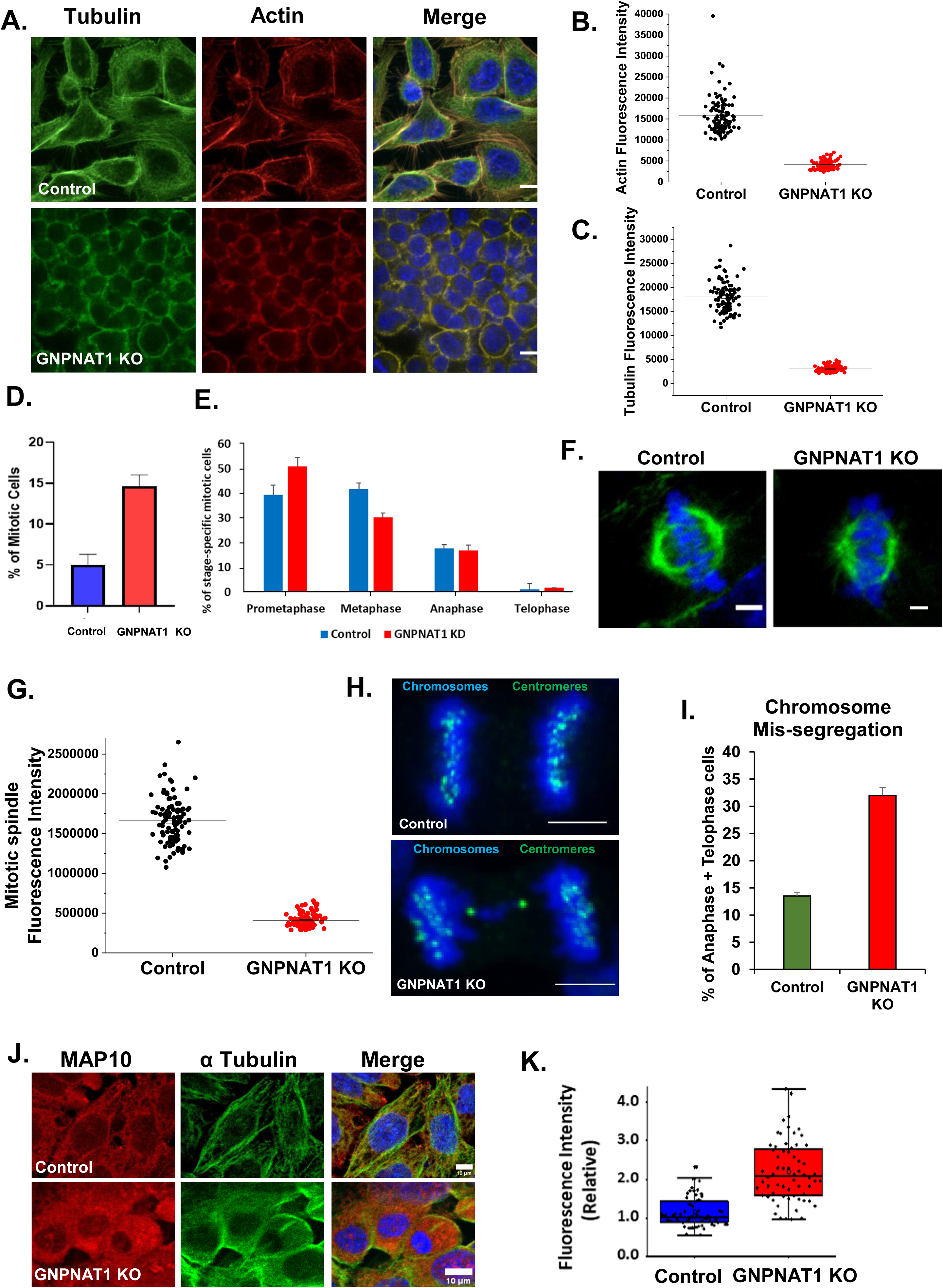
HBP depletion disrupts both actin as well as the microtubule cytoskeleton and increases mitotic frequency. (A) Immunostaining of the actin and microtubule cytoskeleton in control and GNPNAT1 KO 22Rv1 cells. Scale bars, 10 µm. (B-C) Quantification of intensity of individual actin (B) and microtubule (C) filaments after immunostaining. (D) Mitotic cell count of control and GNPNAT1 KO cells, represented as the Mitotic Index (number of mitotic cells/100 total cells). (E) Stage specific mitotic cell count of control and GNPNAT1 KO cells. (F) Immunofluorescence staining of mitotic spindles in control and GNPNAT1 KO cells. Scale bar, 5 µm. (G) Quantification of total microtubule intensity from F. (H) Control and GNPNAT1 KO cells were immuno-stained with a centromere marker and the chromosomes marked using DAPI. Scale bar, 5 µm. (I) Quantification of the frequency of chromosome mis-segregation events from H. (J) Co-staining of MAP10 and microtubules in 22Rv1 control and KO cells. Scale bars, 10 µm. (K) Quantification of MAP10 fluorescence intensity.

Considering the importance of the HBP pathway, we expect significant metabolite changes downstream of the pathway, which may have an impact on the altered cellular behavior observed following HBP inhibition. Of the 288 metabolites examined in our metabolomics studies, we found that 66 had significantly altered levels (Figure 1F). We observed that consistent with our transcriptome analysis, metabolomics analyses additionally revealed how HBP affected multiple other pathways. The heatmap (Figure 1F) shows the significant differences in the metabolites found in GNPNAT1 KO cells as compared to control 22Rv1 cells. These findings indeed suggest that, as expected, our GNPNAT1 KO approach induced substantial alteration to the HBP and various other HBP-related metabolic pathways.

Figure 1E indicated that various pathways were being impacted based on the observed changes in metabolite levels (Supplementary Figure S2F). Most of these pathways are primarily related to the metabolism of amino acids, which regulates cell growth, adhesion, homeostasis, and other aspects of cell signaling. The implications of these metabolite changes on cellular properties prompted us to investigate this further. Given the association between the HBP and protein glycosylation, we conducted a glycoproteomic analysis to explore the impact of HBP on glycosylation. Our analysis revealed that glycosylation of multiple cellular components was altered (Supplementary Figure S2G), which in turn is known to influence various cellular functions (Figure 1I). Since glycosylation is known to control the stability of cellular proteins, we carried out proteomics analyses to assess the alterations in protein levels. These studies suggest that the levels of several 100’s of proteins were differentially altered in GNPNAT1 KO cells as compared to controls (Supplementary Figure S2H).

### HBP inhibition causes defective mitosis

As HBP inhibition has been shown to promote cell proliferation and based on the results from our transcriptomic and metabolomic analysis, we sought to understand more about the downstream genes that HBP pathways regulate to alter the mitotic characteristics of human cells. The total number of proliferating mitotic cells increased significantly in GNPNAT1 KO cells (Figure 2D). Additionally, we classified mitotic cells according to their stages and noticed that there were many more prometaphase cells and fewer metaphase cells after GNPNAT1 KO (Figure 2E). We observed a similar trend for control and GNPNAT1 KO RWPE1 cell lines (Supplementary Figure 3A-B). Further, we found that the structure of the mitotic spindle was substantially altered in GNPNAT1 KO cells as compared to control 22Rv1 cells. We observed a significant reduction in spindle size (Figure 2F; Supplementary figure 3C) with our quantification demonstrating that (there was an approximately three-fold decrease in spindle microtubule intensity (Figure 2G). Spindle disorganization is known to cause chromosome missegregation. These missegregations in turn can drive cell proliferation by promoting aneuploidy, where cells gain an abnormal number of chromosomes. In our study, we observed a threefold increase in chromosomal missegregation in GNPNAT1 knockout cells compared to controls (Figure 2H-I).

We identified that MAP10 is a major potential target downstream of the HBP that showed an enrichment of many mitotic genes based on our previous gene-expression data and mitotic results. The assembly and stability of microtubules are regulated by microtubule-associated proteins (MAPs)(17) and MAPs have been shown to play a critical role in microtubule reorganization of cells(18). MAP10 is a novel microtubule-associated protein that has been reported to stabilize microtubules, and is essential for proper cell division, as its depletion leads to cytokinesis failure and polyploidy. In this study, we observe that, in comparison to control 22Rv1 cells, MAP10 was considerably upregulated in GNPNAT1 KO cells (Figure 2J and 2K). Since MAP10 upregulation after GNPNAT1 inhibition is associated with a reduction in the intensity of microtubule filaments, we believe that MAP10 is possibly a negative regulator of microtubule dynamics. Further studies are required in the future will determine if effects on microtubule cytoskeleton in GNPNAT1 inhibited cells is indeed mediated by alterations in MAP10 function.

### HBP depletion promotes cell proliferation by altering various cell signaling pathways

While the mechanism by which the HBP integrated into various gene expression programs is largely unexplored, this study focuses on the implications of some transcriptional programs that have been established to some extent. Previous studies have indicated that the PI3K/AKT pathway is activated in GNPNAT1 inhibited cells(8). In this study, our results confirmed this finding by demonstrating that when HBP was inhibited, AKT was upregulated in both its total and phosphorylated forms (Figure 3A). Moreover, mTORC1/2 pathway and PKC pathway which are downstream to AKT were also elevated in GNPNAT1 KO cells (Figure 3B-C). Thus, HBP-activated AKT signaling presents a potential target for controlling CRPC. We treated CRPC cells with the AKT inhibitor MK2206 and observed a greater reduction in cell proliferation in GNPNAT1 KO cells (Figure 3D-E). This effect was further confirmed by the rescue of proliferation upon treatment with UDP-GlcNAc (Figure 3F-G). Furthermore, our results confirmed that inhibiting GNPNAT1 phosphorylated and activated FOXO3a, specifically within the nucleus (Figure 4A). Thus, we speculate that FOXO3a is activated by AKT phosphorylation, leading to increased cell proliferation (Figure 3H).

**Figure 3.**
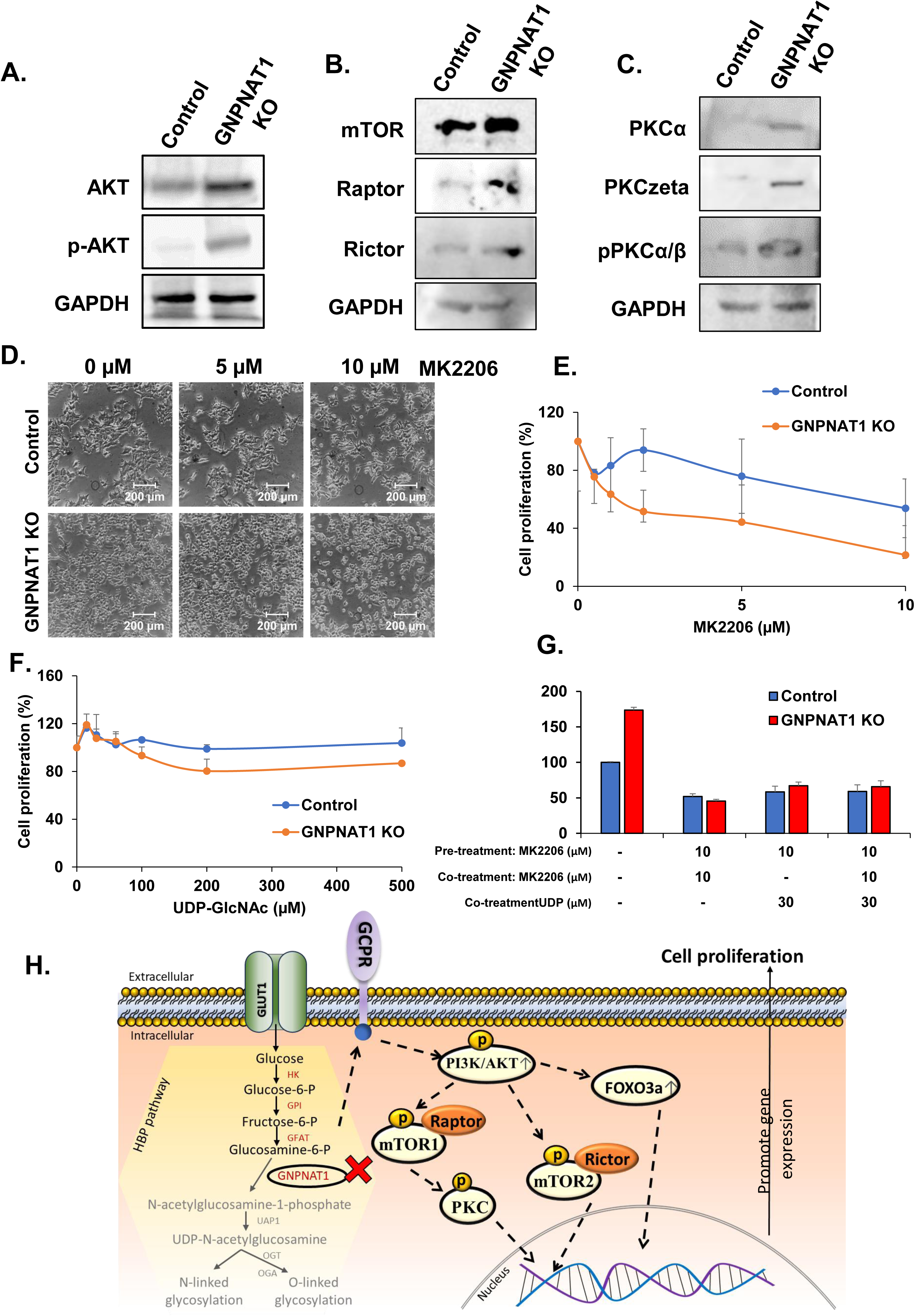
HBP depletion promotes cell proliferation by altering AKT-associated signaling pathways. **(A-C)** Control and GNPNAT1 KO 22Rv1 cell lysate samples were Western blot-analyzed for the levels of AKT and its phosphorylated form p-AKT (A) as well as AKT downstream signaling pathways including components of the mTOR pathway (B), and the PKC pathway (C). (D-E) Microscopic images (D) and cell proliferation assay (E) after treating control and GNPNAT1 KO with the AKT inhibitor, MK2206 after 48 h. Scale bars, 200 µm. F) Cell proliferation assay after treatment with UDP-GlcNAc after 48 h. (G) Cell proliferation assay after pre-treatment with 10 µM MK2206 followed by co-treatment with 10 µM MK2206 and 30 µM UDP-GlcNAc or UDP-GlcNAc alone. (H) Schematic diagram of the possible mechanism for increased cell proliferation in GNPNAT1 KO cells, via the alteration of AKT and its downstream signaling pathways.

**Figure 4.**
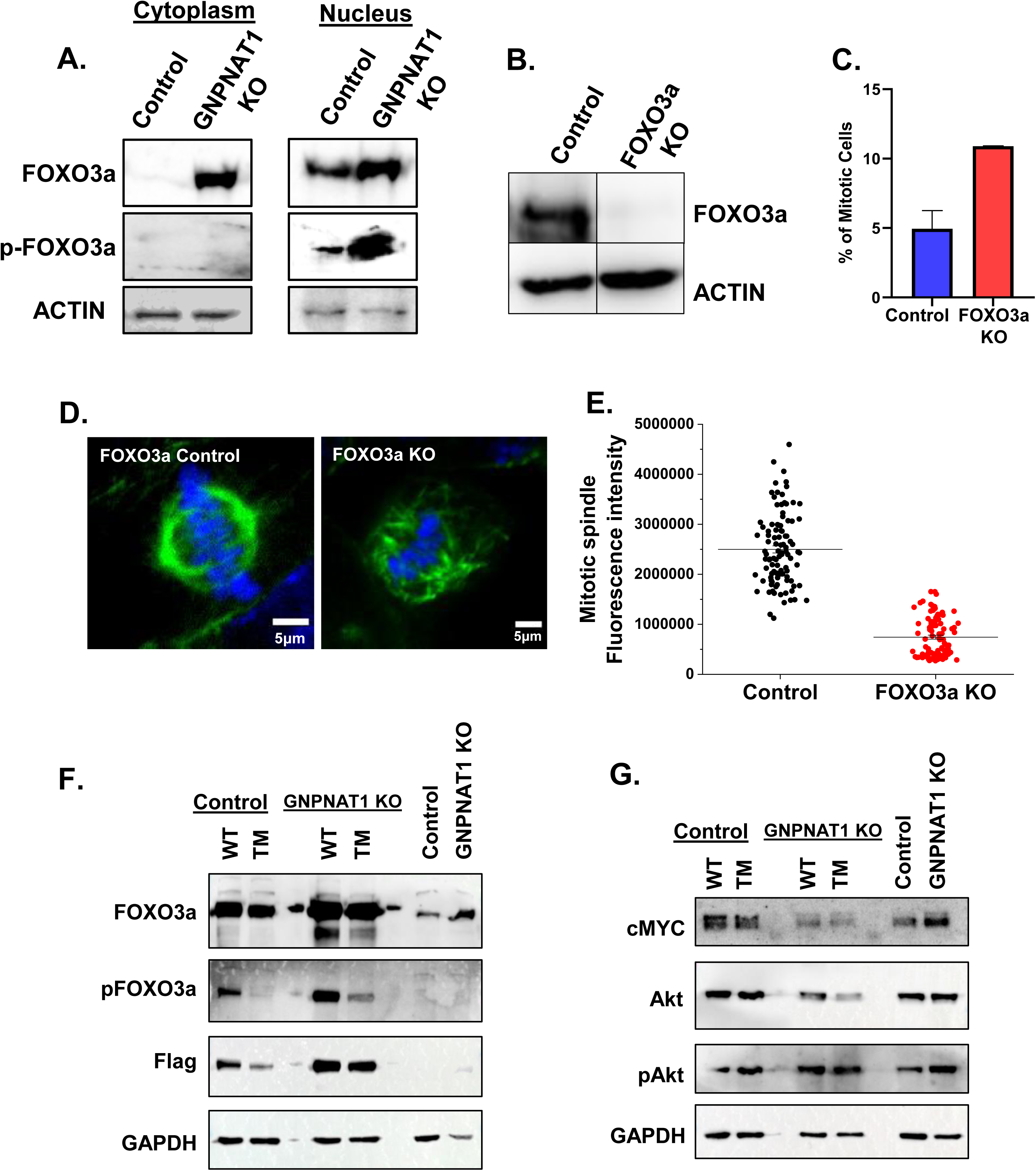
HBP depletion causes an increase in levels of transcription factor, FOXO3a. (A) Cytoplasmic and nuclear levels of FOXO3a and p-FOXO3a in control and GNPNAT1 KO cells assesses by Western Blotting. (B) Western blot confirming CRISPR CAS9-mediated knockout of FOXO3a in 22Rv1 cells. (C) FOXO3a KO samples were analyzed for the number of mitotic cells/100 total cells (Mitotic Index) relative to control cells. (D) The same cells were immuno-stained for microtubules, with the chromosomes counterstained using DAPI. Scale bars, 5 µm. (E) Total intensity of the mitotic spindle microtubules was quantified in controls vs FOXO3a KO cells. (F-G) Expression levels of FOXO3a, p-FOXO3a, and FLAG tag (F) as well as cMYC, AKT, and p-AKT (G) in cells expressing WT and TM (non-phosphorylatable mutant) FOXO3a.

FOXO3a is a transcription factor that regulates a variety of processes including cell survival, proliferation, and apoptosis(19). Following AKT-mediated phosphorylation (at Thr 32, Ser 253, and Ser 315), Foxo3a is exported from the nucleus, where it associates with 14-3-3 proteins prior to its retention in the cytoplasm(20). We created FOXO3a KO in the 22Rv1 CRPC cells, which we confirmed by Western blot(Figure 4B). This experiment was designed primarily for comparison with the GNPNAT1 KO studies. Our findings demonstrate that, when compared to control 22Rv1 cells, the FOXO3a knockout causes a doubling of mitotic cells (Figure 4C). Further, the deletion of FOXO3a decreased the intensity of mitotic spindle microtubules by a factor of more than three. These results suggest that the loss of FOXO3a phenocopies that of GNPNAT1 KO (Figure 4D-E). On the other hand, overexpression of FOXO3a WT as well as FOXO3a non-phosphorylatable mutant (TM) (Figure 4F) causes a reduction in expression levels of AKT as well as cMYC in GNPNAT1 KO cells as compared to control cells (Figure 4G). The physiological basis of this observation is not clear at this point.

### HBP depletion alters the cell adhesion and migration properties

Upon closely examining the morphology of the cells, we observed that the GNPNAT1 KO 22Rv1 cells were more rounded or unusually elongated and occupied less area than the control 22Rv1 cells (Figure 5A). In order to investigate the effects of GNPNAT1 knockout on cell adhesion to the substratum, we performed a cell adhesion assay in which we monitored the spreading of GNPNAT1 KO and control 22Rv1 cells on fibronectin-coated glass-bottom dishes. Time-lapse imaging was used to record the cell morphology, with a 15-min frame interval (Figure 5B-C). Our studies indicated that the 22Rv1 KO cells spread at a much slower rate as compared to the KO cells, suggesting that the cell adhesion had been impaired in the KO cells (Figure 5B-C). Based on the results from wound healing experiments of 22Rv1 control and GNPNAT1 KO cells, we observed that GNPNAT1 KO cells demonstrated enhanced cell migration properties (Figure 5D-F). The same pattern was observed in control and GNPNAT1 RWPE1 cell lines (Supplementary Figure S4A-C). Since the wound healing assay only represents cohort cell migration, we performed trans-well cell invasion/migration assay to confirm an effect on individual cell migration. Based on this assay, we observed that cells migrate and invade more in GNPNAT1 KO cells in comparison to control cells (Figure 5G).

**Figure 5.**
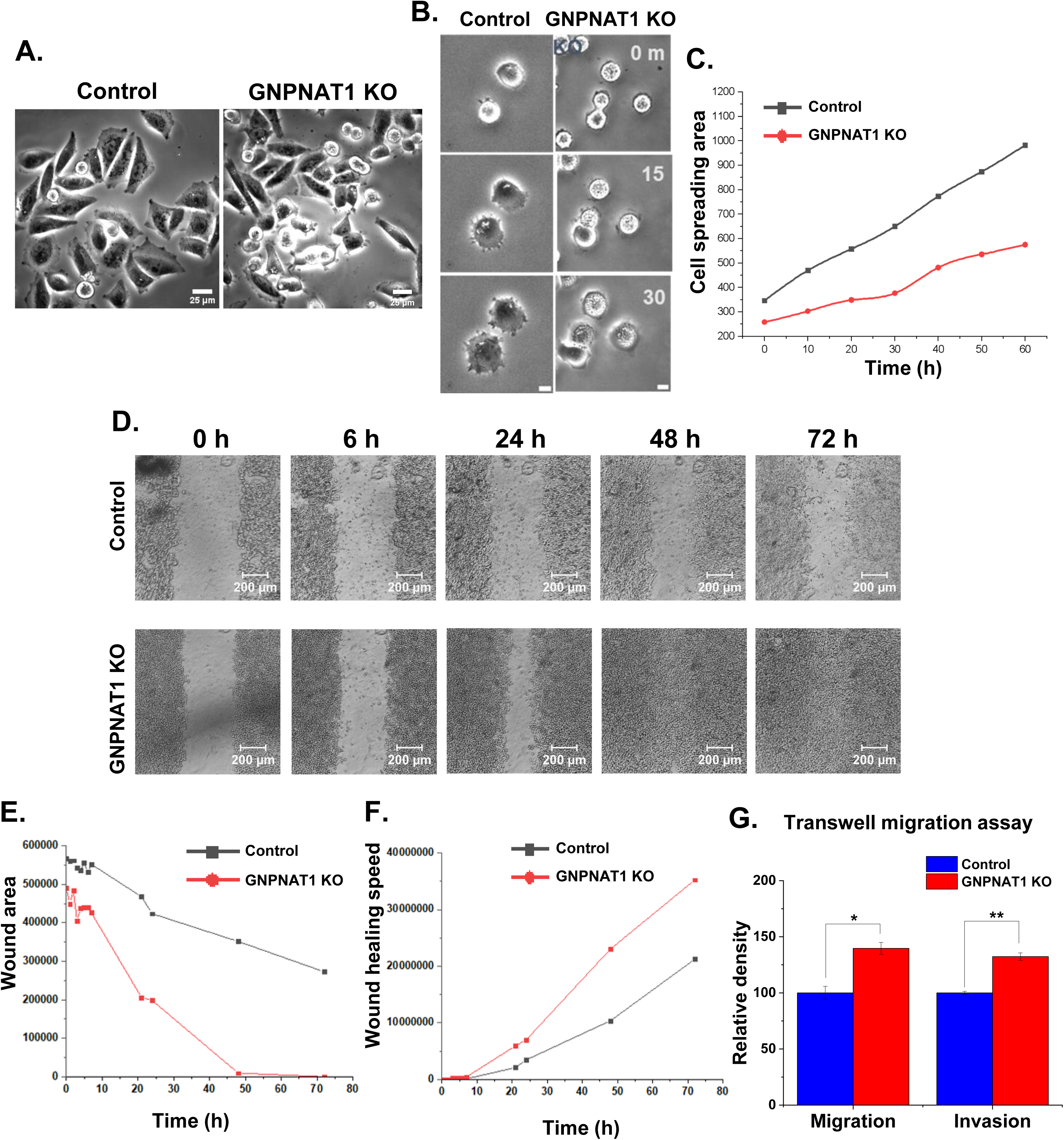
HBP depletion impacts cell adhesion to the sub-stratum and also accelerates cell migration. (A) Confocal microscopic images of control and GNPNAT1 KO 22Rv1 cells, showing cell morphology. Scale bars, 25 µm. (B-C) Microscopic images (B) and quantification of cell spreading (C) performed on fibronectin-coated glass surface. Scale bars, 25 µm. (D-F) Microscopic images (D) and graph showing wound area (E) and wound healing speed (F) observed by the wound healing assay. Scale bars, 200 µm. (G) Transwell migration and invasion assay of control and GNPNAT1 KO 22Rv1 cells.

Ephrin and Eph receptors are overexpressed in a variety of human tumors, as shown by multiple studies(21,22). On the other hand, tumor development has been associated with both their up- and down-regulation, and both ephrin ligands and Eph receptors have the ability to either promote or inhibit the growth of tumors(23,24). We observed that EphB6 is downregulated in GNPNAT1 KD cells as initially suggested by our RNA sequencing data (Figure 6A-B). This is what we believe, is a contributing factor in the reduction of cell adhesion properties. Further, we observed that several other Eph receptors were also downregulated in addition to EphB6 after GNPNAT1 KO as demonstrated by the RTK proteome array (Figure 6C-D), which is consistent with the transcriptomics results.

**Figure 6.**
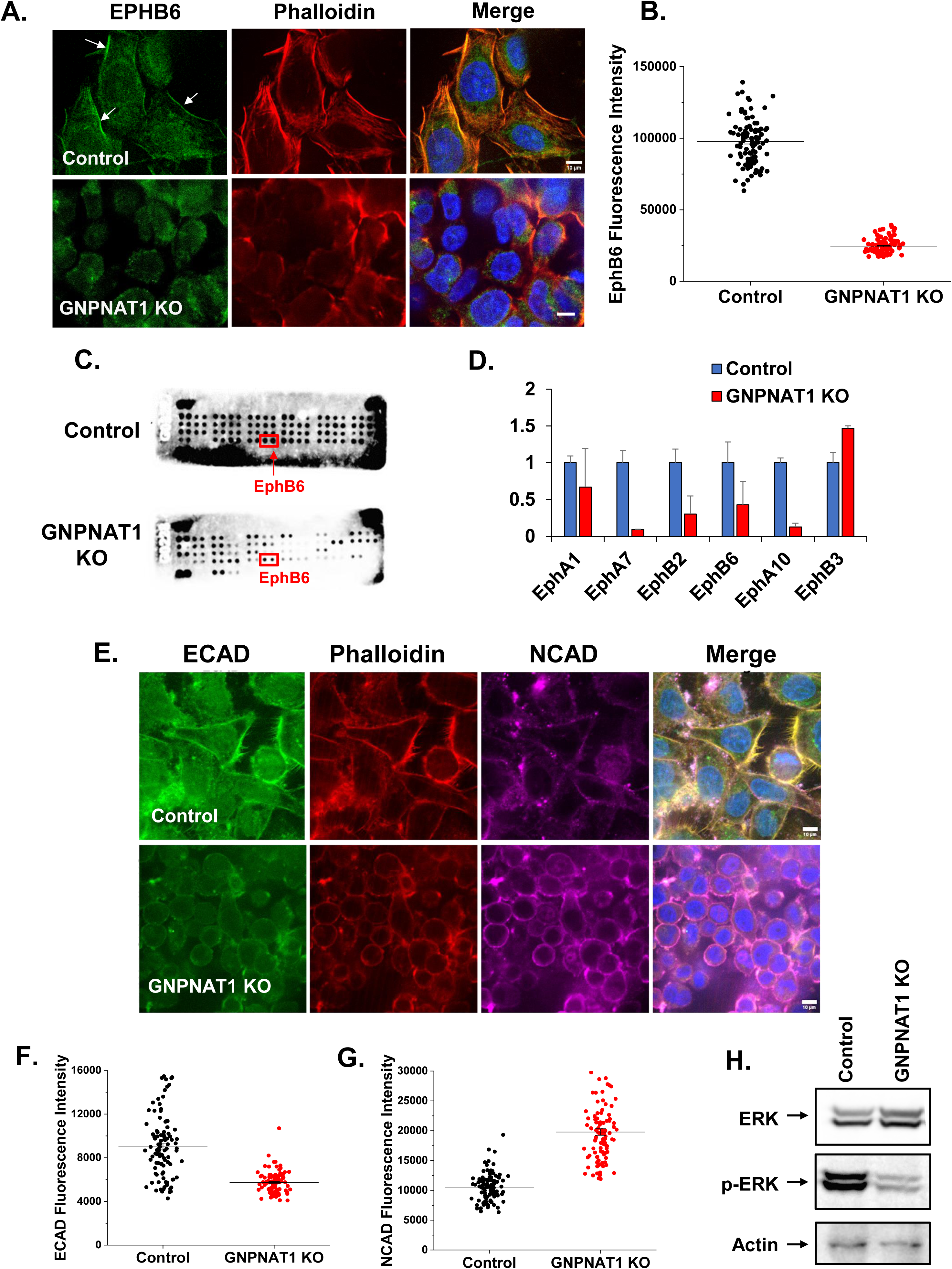
HBP depletion reduces EphB6 ERK signaling and promotes Epithelial-to-Mesenchymal Transition (EMT). (A) EphB6 immunofluorescence staining with DAPI in control and GNPNAT1 KO 22Rv1 cells. Arrows indicate brighter EphB6 staining at the cell periphery. Scale bars, 10 µm. (B) Quantification of EphB6 levels at the cell periphery from A. (C-D) Quantification of protein expression level of multiple Ephrins using Human RTK proteome array. (E) ECAD, NCAD and Phalloidin-actin immunofluorescence staining with DAPI in control and GNPNAT1 KO 22Rv1 cells. Scale bars, 10 µm. (F-G) Quantification of ECAD and NCAD levels at the cell periphery from E. (H) Expression levels of total and phosphorylated form of ERK in control and GNPNAT1 KO 22Rv1 cells as indicated.

Our previous results had led us to insinuate that epithelial-to-mesenchymal transition (EMT) is crucial to the observed cellular phenotype as we had identified notable changes in the extracellular matrix (ECM) assembly and disassembly pathways. Figure 6E-G illustrates that the E-Cadherin levels were lower in GNPNAT1 KO cells while NCAD levels were elevated. Overall, the loss of epithelial cadherin (E-cadherin) and the replacement of N-cadherin is a characteristic of epithelial-to-mesenchymal transition. The EMT phenotype in CRPC cells is indeed influenced by GNPNAT1 downregulation, as evidenced by the immunofluorescence results for E- and N-cadherin. We also find that the levels of phosphorylated ERK was reduced in GNPNAT1 inhibited cells (Figure 6H). Based on these results, we speculate that downregulation of EphB6 activated the MAPK/ERK signaling pathway to alter cell surface cadherins and drive the loss of adhesion, which in turn might facilitate cell migration (Figure 7E). As initially suggested by our RNA seq results, our immunofluorescence staining experiments confirmed that the expression of the RND3 is reduced in GNPNAT1-inhibited cells. These experiments also showed that RhoA levels increases in the same condition (Figure 7A-D). It is known that downregulation of RND3 leads to increased RhoA activity which enhances cell migration, invasion, and other mesenchymal behaviors through epithelial to mesenchymal transition(25–28). Moreover, other Rho members such as RhoB and RhoC were found to be reduced after GNPNAT1 inhibition (Supplementary Figure 5A-D).

**Figure 7.**
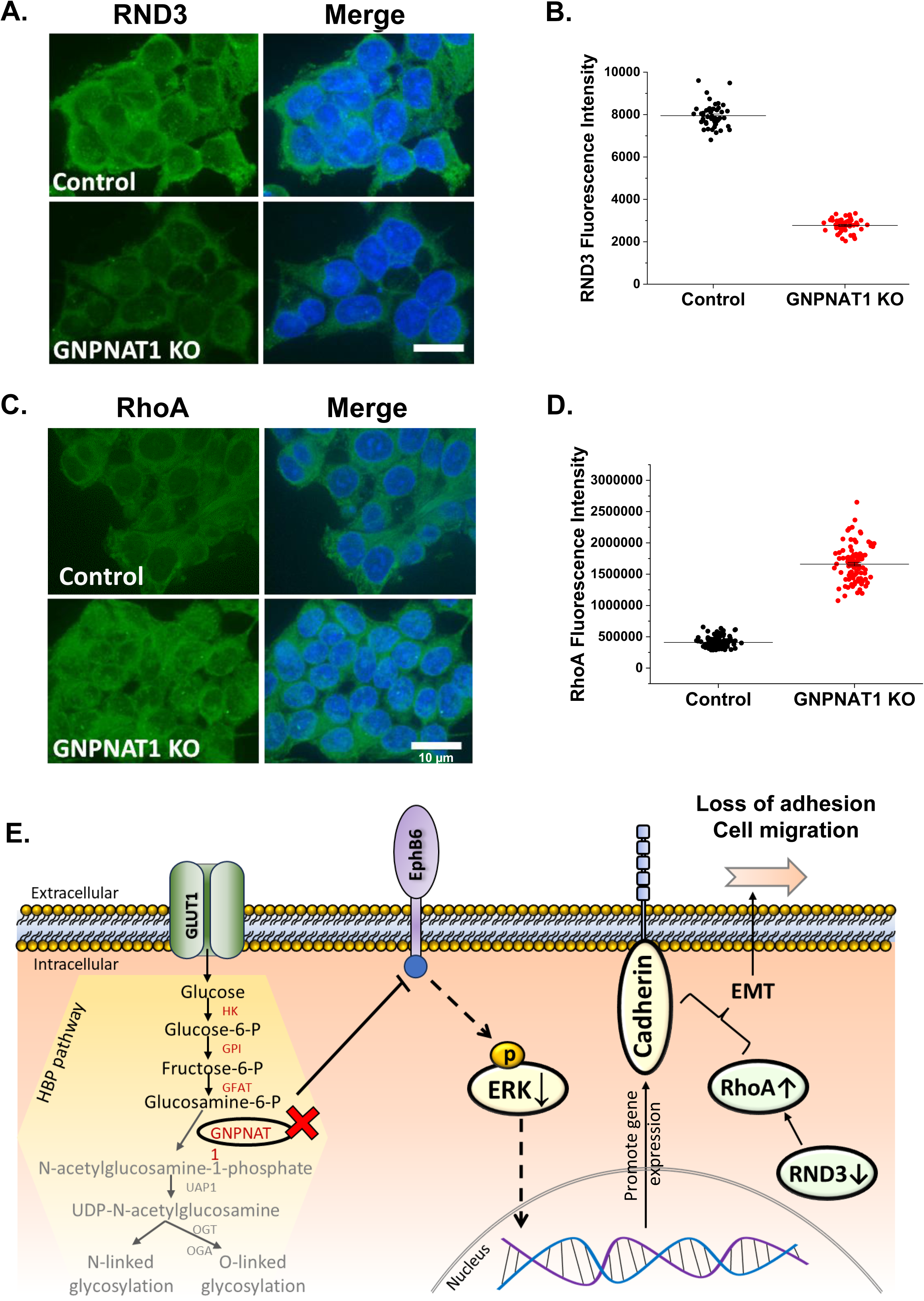
HBP depletion alters RND3/RhoA signaling. (A) RND3 immunofluorescence staining with DAPI in control and GNPNAT1 KO 22Rv1 cells. (B) Quantification of RND3 cytoplasmic levels from A. (C) RhoA immunofluorescence staining with DAPI in control and GNPNAT1 KO 22Rv1 cells. (D) Quantification of RhoA cytoplasmic levels from C. Scale bars, 10 µm. (E) Schematic diagram of the possible mechanism of how HBP KO impacts the loss of cell adhesion and promote cell migration.

## Discussion

Previous work had combined gene-expression, mouse in vivo studies and metabolomics analysis on benign and prostate cancer tissues(8,29). Compilation of gene-expression and metabolomic data showed overall enrichment of five pathways including riboflavin metabolism, biotin metabolism, amino sugar metabolism (HBP), valine, leucine, and isoleucine biosynthesis, and cysteine metabolism. Among various pathways studied, the HBP was found to have the highest level of interactions, which shows that this pathway is highly significant within biological networks and is central to many cellular processes. This analysis was based on data from the KEGG (Kyoto Encyclopedia of Genes and Genomes) database, which maps biological systems. The finding highlights the importance of the HBP in regulating key biological functions due to its extensive involvement in various cellular interactions.(8).

In HBP, glucose is transported into the cell and gets phosphorylated to form glucose-6-phosphate, which gets isomerized to fructose-6-phosphate. Amidation of fructose-6-phosphate with glutamine produce glucosamine-6-phosphate(30). GNPNAT1 catalyzes the transfer of an acetyl group from acetyl-CoA to glucosamine-6-phosphate, forming N-acetylglucosamine-6-phosphate (GluNAc-6-P). The enzyme phosphoglucomutase catalyzes the conversion of GluNAc-6-P to N-acetylglucosamine-1-phosphate (GluNAc-1-P). Then, uridyltransferase transfers a uridine monophosphate (UMP) group from UTP (uridine triphosphate) to GluNAc-1-P, resulting in the formation of UDP-N-acetylglucosamine (UDP-GluNAc), a molecule crucial for glycosylation. Glycosylation is essential for modifying proteins and plays a role in various cellular processes, including signal transduction and protein stability(11,30,31).

Several studies have observed the upregulation of gene-expression profiles of HBP genes such as GNPNAT1, UAP1, and UDP-GluNAc in localized prostate cancer tissue(14,29,32,33). However, the expression profiles of these gene were significantly reduced in metastatic or CRPC tumor tissue. Further, GNPNAT1 knockdown leads to increased cell proliferation and metastasis of 22Rv1 and LNCaP CRPC cell lines. Meanwhile, over-expression of GNPNAT1 in 22Rv1 cells results in the significant increase in cell proliferation 6 days post transfection(8,14).

The exact mechanism of elevated cell proliferation and metastasis of HBP-inhibited cells are unknown. Thus, we carried out the knockout of the GNPNAT1 gene in 22Rv1 human CRPC cell line using CRISP-Cas9 to better understand the functional implications of this pathway to cellular behavior. After GNPNAT1 knockout, cell morphology undergoes significant alterations with an increase in cell proliferation. One reason for this abnormal cell proliferation is elevated mitotic cell numbers in GNPNAT1 KO cells. It is not clear at this point if this is brought about by an increased number of cells entering mitosis and/or delayed progression of cells through mitosis, thus causing an increase in mitotic cells at any given time. Further, we find that the intensity of spindle microtubules and the size of the spindles as such is reduced in the GNPNAT1 KO cells in comparison to control 22Rv1 cells. While there is limited physiological evidence to support this notion, it is also possible that in GNPNAT1 KO cells, the smaller mitotic spindle might contribute to faster cell division by accelerating the separation of chromosomes and completing mitosis more quickly thus leading to an overall increase in the rate of cell proliferation, but the accuracy of chromosome segregation could be compromised, as we see increased chromosome mis-segregation after GNPNAT1 KO.

As with interphase cells, microtubules also undergo dynamic changes, including polymerization and depolymerization during mitosis, which in turn controls the integrity of the mitotic spindle(34). Certain proteins that bind to the microtubules, referred to as Microtubule-Associated Proteins (MAPs) are involved in the regulation of microtubule dynamics. These MAPs are important to maintain the architecture of the mitotic spindle in dividing cells, which in turn is critical for the accurate alignment and segregation of chromosomes(35–37). MAP10 is a newly identified MAP that has been reported to enhance microtubule stability and play a role in ensuring successful cytokinesis and preventing polyploidy(38). However, a thorough functional characterization of MAP10 has not been carried out so far. In our studies, we find that MAP10 expression is elevated after the loss of GNPNAT1. However, we also find that the intensity of microtubules in interphase and mitotic cells are reduced, and that mitotic spindle morphology is altered in this scenario. Further work will determine whether these defective mitotic phenotypes observed after HBP inhibition is indeed mediated via the interference of MAP10 function.

A high incidence of chromosome mis segregation was observed in GNPNAT1 KO cells. This leads to aneuploidy, a condition where cells have abnormal number of chromosomes(39). While aneuploidy is often associated with cell dysfunction and disease, in some cases, it has been observed to increased cell proliferation. Aneuploid cells may experience growth deficiencies due to the imbalance in gene dosage caused by the abnormal chromosome number(40). To compensate for this, cells might enter the cell cycle more frequently, in an attempt to overcome the challenges posed by aneuploidy, thus leading to the observed increase in proliferation. In addition, aneuploid cells may activate certain adaptive responses that enhance cell survival and proliferation. These responses could include changes in gene expression patterns, activation of specific signaling pathways, or alterations in the cell cycle checkpoints(40–42).

Furthermore, the cytoskeleton, including both actin and microtubules, reorganizes to support the mitosis, cell proliferation, and migration process(43). Actin filaments facilitate the formation of cellular protrusions, enabling movement, while microtubules provide structural support and guide intracellular trafficking during migration. Actin and microtubules are the dynamic players orchestrating the intricate processes of cell proliferation and migration, ensuring the proper division and movement of cells(44,45). The disorganization as well as reduced intensity of both these cytoskeletal structures were observed in GNPNAT1 KO cells. Loss of these cytoskeletal structures affect the dynamic network of intracellular structures that provide structural support to maintain cell shape, cell motility, as well as cell division and also influence signaling pathways.

It is known that PI3k/AKT pathway is activated when GNPNAT1 is inhibited(8). The signaling between AKT and FOXO3a is a well-studied as an intricate regulatory pathway and a crucial signaling axis involved in regulating cell survival as well as proliferation(46). In GNPNAT1 KO cells, AKT is phosphorylated leading to its activation followed by phosphorylation of FOXO3a. We predict that the increased FOXO3a is responsible for controlling cytoskeletal genes and promoting accelerated cell division.

AKT regulates numerous downstream effectors by phosphorylating and modulating their activities. mTOR, or mammalian target of rapamycin, is a crucial protein kinase that forms two distinct complexes, mTORC1 and mTORC2(47,48). mTORC1 primarily regulates cell growth and metabolism, while mTORC2 regulates cytoskeletal organization and cell survival. Both mTOR complexes are sensitive to various growth factors, nutrients, and cellular energy levels. Activation of mTORC1 leads to the phosphorylation of several targets involved in translation initiation, ribosome biogenesis, and lipid metabolism, ultimately promoting cell growth. AKT can directly activate mTORC1 through phosphorylation of its inhibitor, TSC2 (tuberous sclerosis complex 2). TSC2 normally inhibits mTORC1 by inhibiting the small GTPase Rheb, which is required for mTORC1 activation. Phosphorylation of TSC2 by AKT relieves this inhibition and allows mTORC1 to be activated. On the other hand, mTORC1 can also indirectly regulate AKT activity. Activation of mTORC1 leads to increased phosphorylation and activation of S6K1, a downstream effector of mTORC1. Activated S6K1 negatively regulates insulin receptor substrate-1 (IRS-1) through phosphorylation, thereby inhibiting the PI3K/AKT signaling pathway. This mechanism provides a negative feedback loop that dampens the AKT signaling pathway when mTORC1 is hyperactive(48–50).

In addition to their direct interactions, both AKT and mTOR can regulate Protein kinase C (PKC) activity. PKC is a family of serine/threonine kinases consisting of multiple isoforms. PKC isoforms have diverse functions and regulate various cellular processes, including signal transduction, cell proliferation, differentiation, apoptosis, and cytoskeletal changes(51). AKT has been shown to phosphorylate and activate certain PKC isoforms, such as PKC-alpha and PKC-zeta, through direct phosphorylation. Activation of PKC by AKT can further promote cell survival and proliferation(52). On the other hand, mTORC1 has been implicated in regulating PKC activity indirectly. mTORC1 signaling can modulate the availability of cellular lipids, which are essential cofactors for the activation of several PKC isoforms. Thus, mTORC1 activity can impact PKC function via lipid metabolism. In general, the relationship between AKT, mTOR, and PKC is complex and involves a network of interactions and feedback loops(53). AKT can directly activate mTORC1 and indirectly regulate PKC activity. mTORC1, in turn, can regulate AKT activity and indirectly influence PKC through lipid metabolism. These interactions and regulations contribute to the coordination of cell growth, metabolism, survival, and other cellular processes(54). HBP inhibited-AKT activation promotes mTOR and PKC favoring for cell proliferation.

Apart from alteration in cell morphology and cell proliferation, recent studies also showed the involvement of HBP in metastasis. Based on our cell spreading assays, GNPNAT1 KO cells were observed to have impaired ability to spread and attach to the plate indicating reduced cell adhesion. This decrease in cell adhesion in HBP depleted cells might be influenced or associated with altered cytoskeletal dynamics or impaired signaling pathways related to adhesion. The results from wound healing and invasion assay indeed demonstrate the enhanced cell migratory and invasive properties of HBP inhibited cells.

Eph receptors and their ephrin ligands play crucial roles in cell communication, migration, and adhesion during development and tissue homeostasis(23). EphB6, specifically, has been associated with multiple cellular functions, including its involvement in cell adhesion and migration processes. HBP-inhibited cells were found to have reduced EphB6 levels. The loss of EphB6 could either lead to the upregulation or activation of other signaling pathways or target its downstream signaling that enhance cell migration while negatively impacting adhesion(24). EphB6, as a member of the Eph receptor family, can activate intracellular signaling pathways, including the MAPK/ERK pathway(21–24). We speculate that the downregulation of EphB6-mediated by HBP inhibition might lead to the suppression of ERK phosphorylation.

Epithelial-to Mesenchymal Transition (EMT) is a biological process where epithelial cells lose their characteristic features, such as cell-cell adhesion and apical-basal polarity(55). This loss of cell-cell adhesion facilitates increased mobility of individual cells, enabling them to migrate independently(56). HBP was associated with EMT where it is accompanied by changes in the expression of various genes and proteins, including downregulation of epithelial marker such as E-cadherin and upregulation of the mesenchymal marker such as N-cadherin. Mesenchymal cells are more motile and can detach from the epithelial layer, allowing them to migrate through the extracellular matrix (ECM) and invade surrounding tissues. EMT is often associated with metastasis by influencing cytoskeletal dynamics and expression of migration-related genes(55,56). In GNPNAT1 KO cells, we observed higher levels of RhoA, and reduced levels of RhoA antagonist, RND3. This could possibly mediate EMT, which in turn drives cytoskeletal reorganization and enhances cell proliferation as well as migration.

Targeting the HBP or its downstream activators such as AKT, FOXO3a, Myc, and RhoA pathways that is implicated in increased cell proliferation and metastasis could be an effective strategy for CRPC treatment. Designing drugs or treatments to inhibit or regulate the function of HBP should disrupt the cancer cells growth and survival mechanisms. Additionally, understanding the specific signaling pathways influenced by the HBP could guide interventions to modulate these pathways in cancer.

## Acknowledgements

We would like to thank Drs. Ajay Abraham and Vipul Shukla (Northwestern University) for their assistance with electroporation. As indicated in the methods, we would like to acknowledge both the NUSeq Core and Proteomics Core facilities at Northwestern University for RNA-sequencing and the Glyco-proteomics analyses respectively. We would like to thank the Beth Israel Deaconess Medical Center-Mass spectrometry Core Facility (Harvard Medical School Teaching Hospital) for Metabolomics analysis. We would also like to acknowledge the Center for Advanced Microscopy & Nikon Imaging Center (Northwestern University) for various help with cell imaging. This work was supported by an R01 grant GM135391 from NIGMS, a SPORE in Prostate Cancer grant P50 CA180995 from NCI and Northwestern University Start-up funds.

## Supplementary figure legends

**Supplementary Figure S1.**
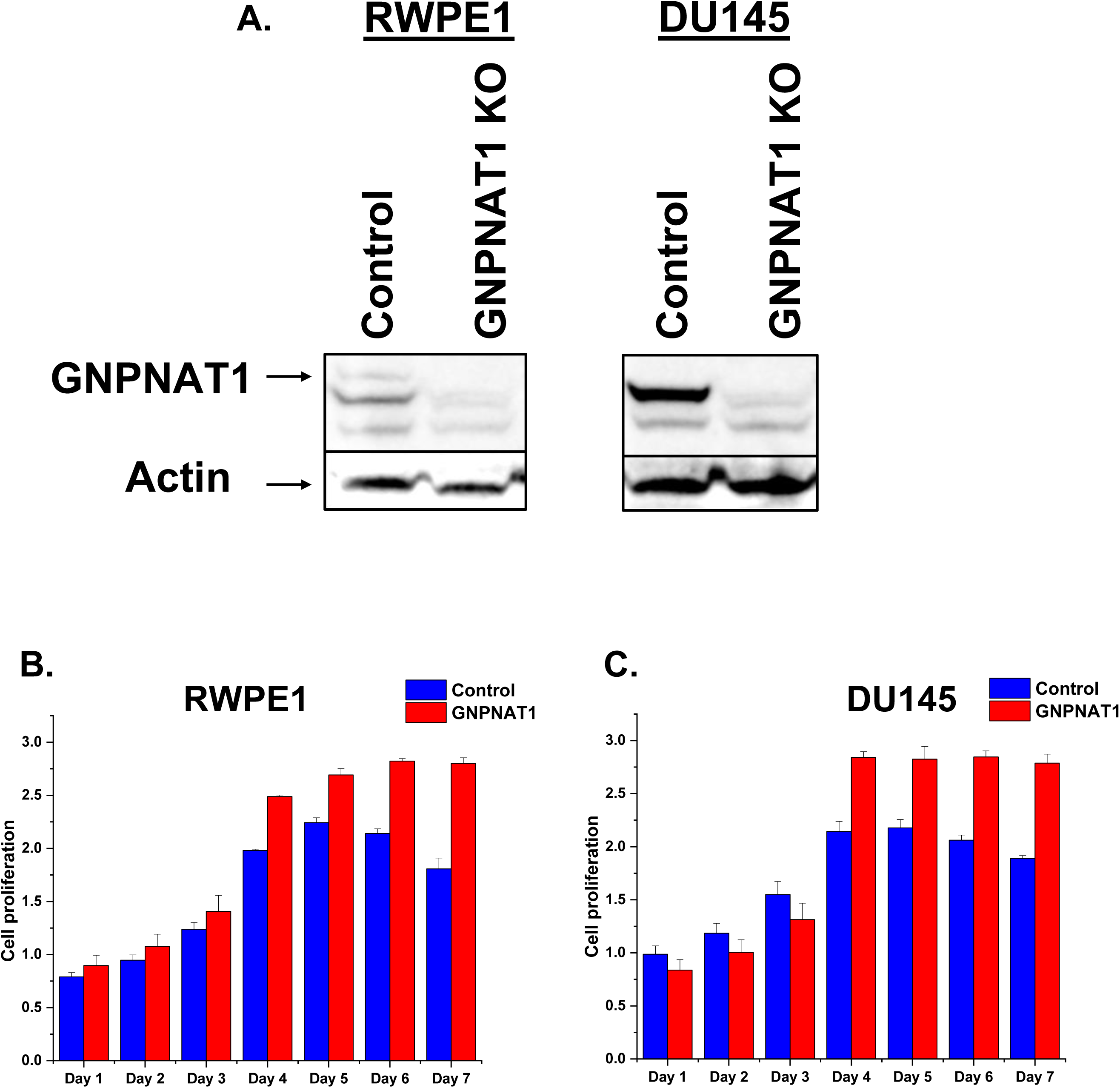
HBP depletion affect the cell proliferation of RWPE and DU145 cell lines. (A) Confirmation of CRISPR CAS9-mediated knockout of GNPNAT1 in RWPE1 and DU145 cells by Western blotting. (B-C) Cell proliferation assay of control and GNPNAT1 KO RWPE1 and DU145 cells at day 1-7.

**Supplementary Figure S2.**
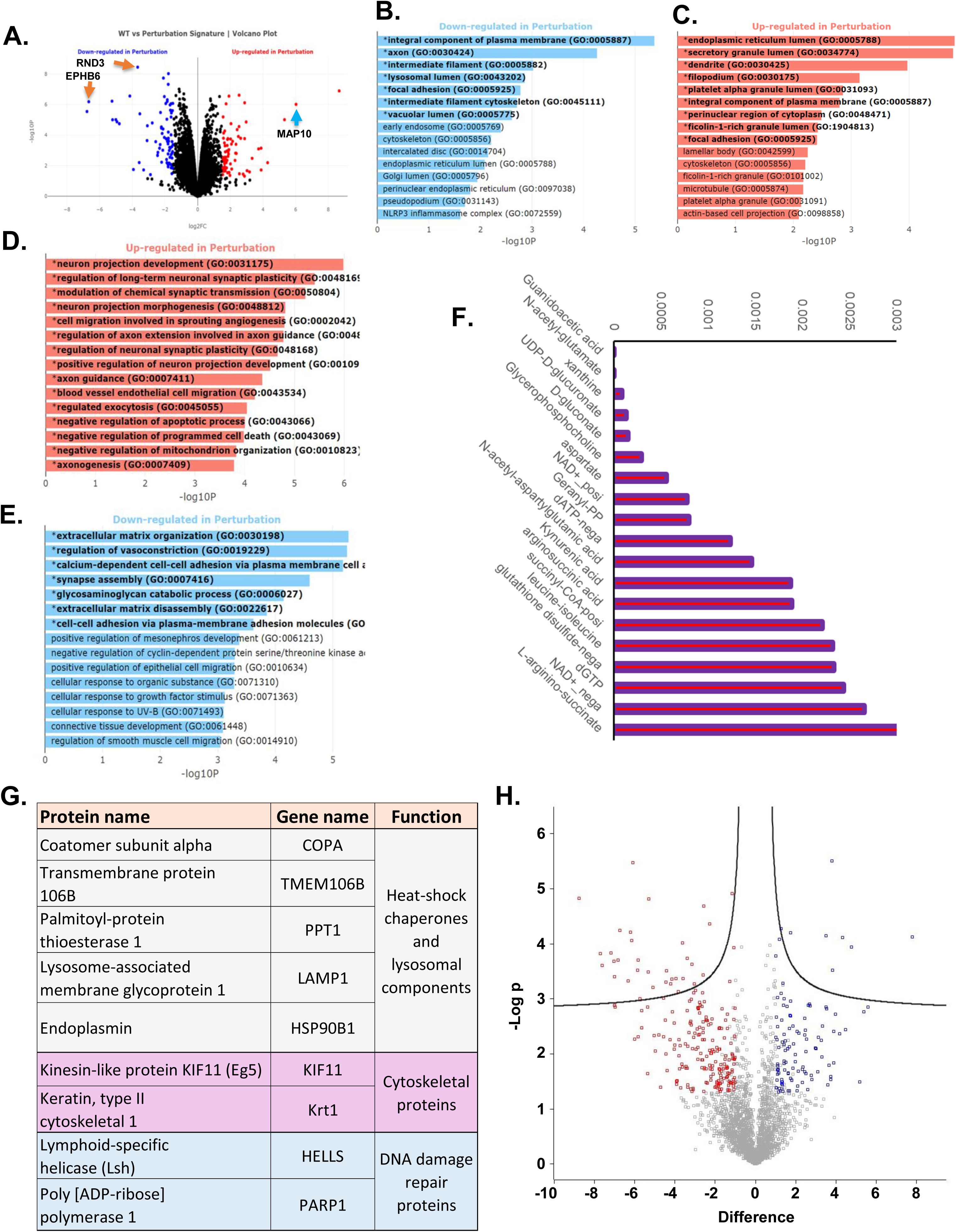
HBP inhibition affects various genes, metabolites, and glycoproteins associated with cell proliferation, adhesion and migration. (A) Volcano plot showing 994 genes being differentially expressed. (B-E) Gene ontology analysis of downregulated (B) and upregulated (C) cellular components as well as upregulated (D) and downregulated (E) biological processes. (F) Metabolomics analysis showing alteration in various metabolites after GNPNAT1 inhibition. (G) A table highlighting nine significant proteins that were shown to be differentially glycosylated. (H) Volcano plot illustrating the differential cellular levels of several 100’s of cellular proteins. Blue spots depict proteins whose levels were elevated while red spots are proteins whose levels were reduced.

**Supplementary Figure S3.**
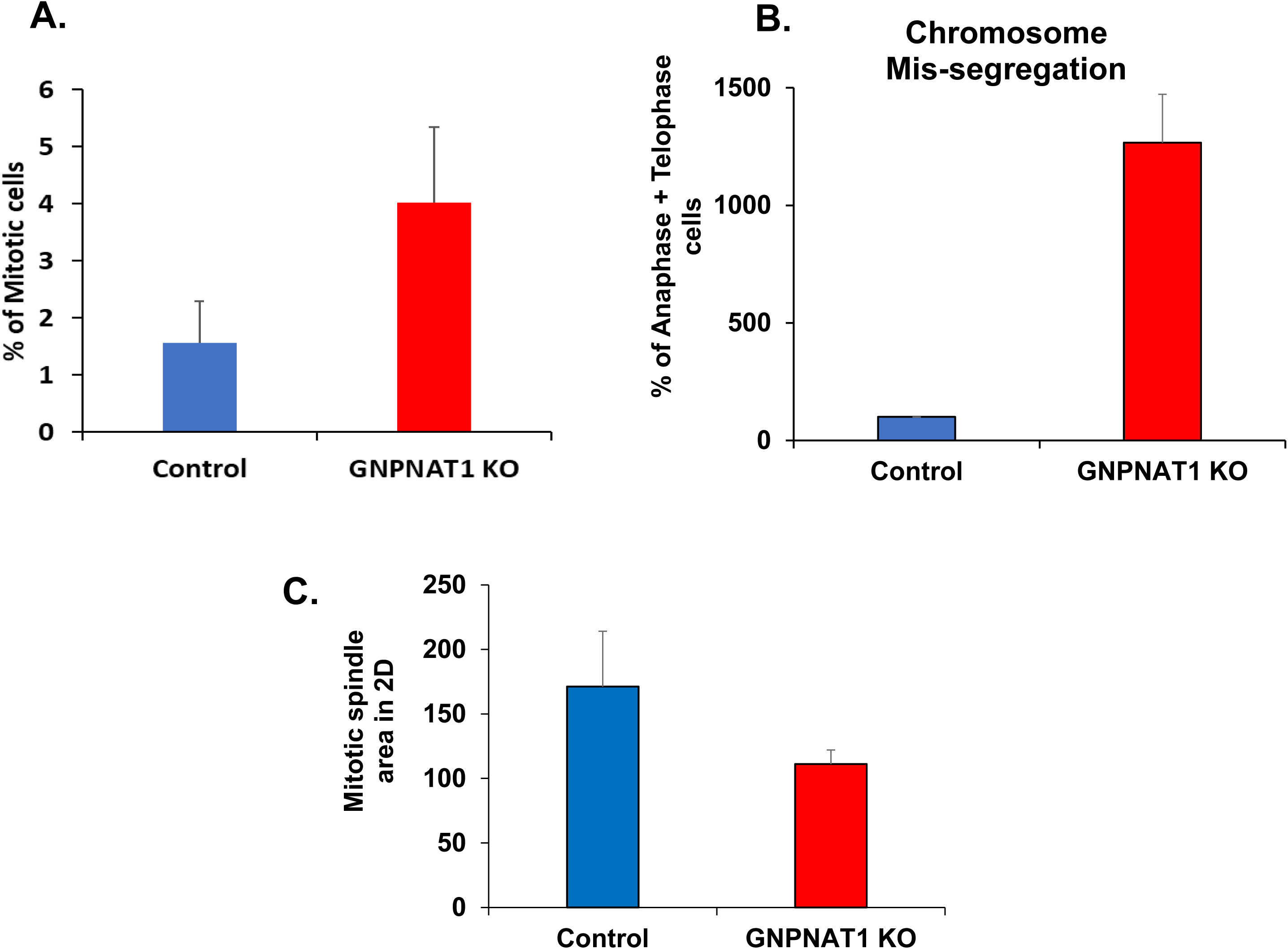
HBP depletion increases the frequency of mitotic cells with chromosome mis-segregation in RWPE cell line. (A) Mitotic cell count of RWPE1 in control and GNPNAT1 KO cells represented as the mitotic index (number of mitotic cells/100 total cells). (B) Analysis of chromosome mis-segregation in control and GNPNAT1 KO RWPE1 cells. (C) Quantification of mitotic spindle area in control and GNPNAT1 KO 22Rv1 cells.

**Supplementary Figure S4:**
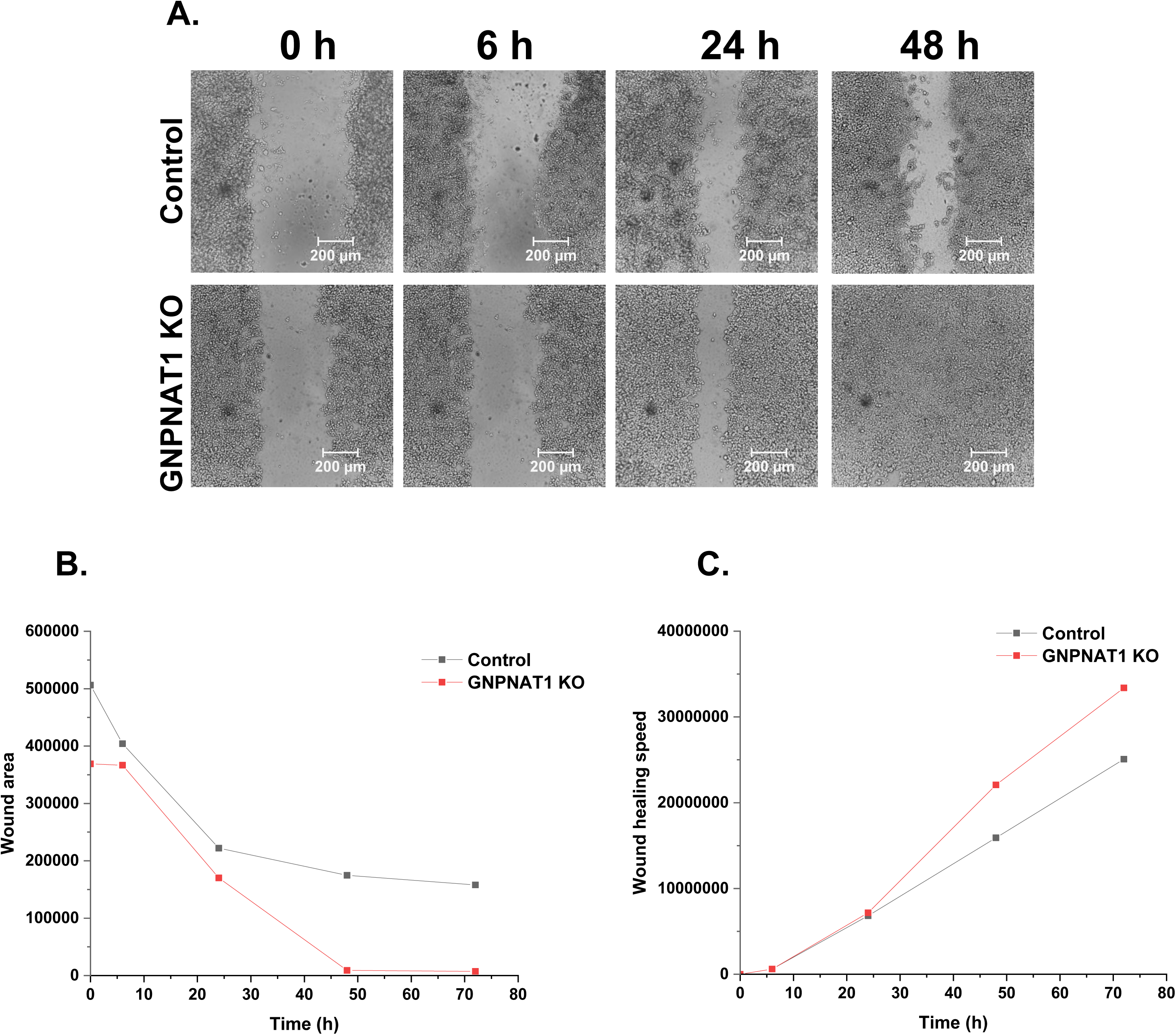
HBP depletion affect the cell adhesion and migration properties of RWPE1 cells. (A) Wound healing assay of RWPE1 control and GNPNAT1 KO cells. Scale bars, 200 µm. (B-C) Quantification of wound healing area (B wound healing speed (C) from A.

**Supplementary Figure S5:**
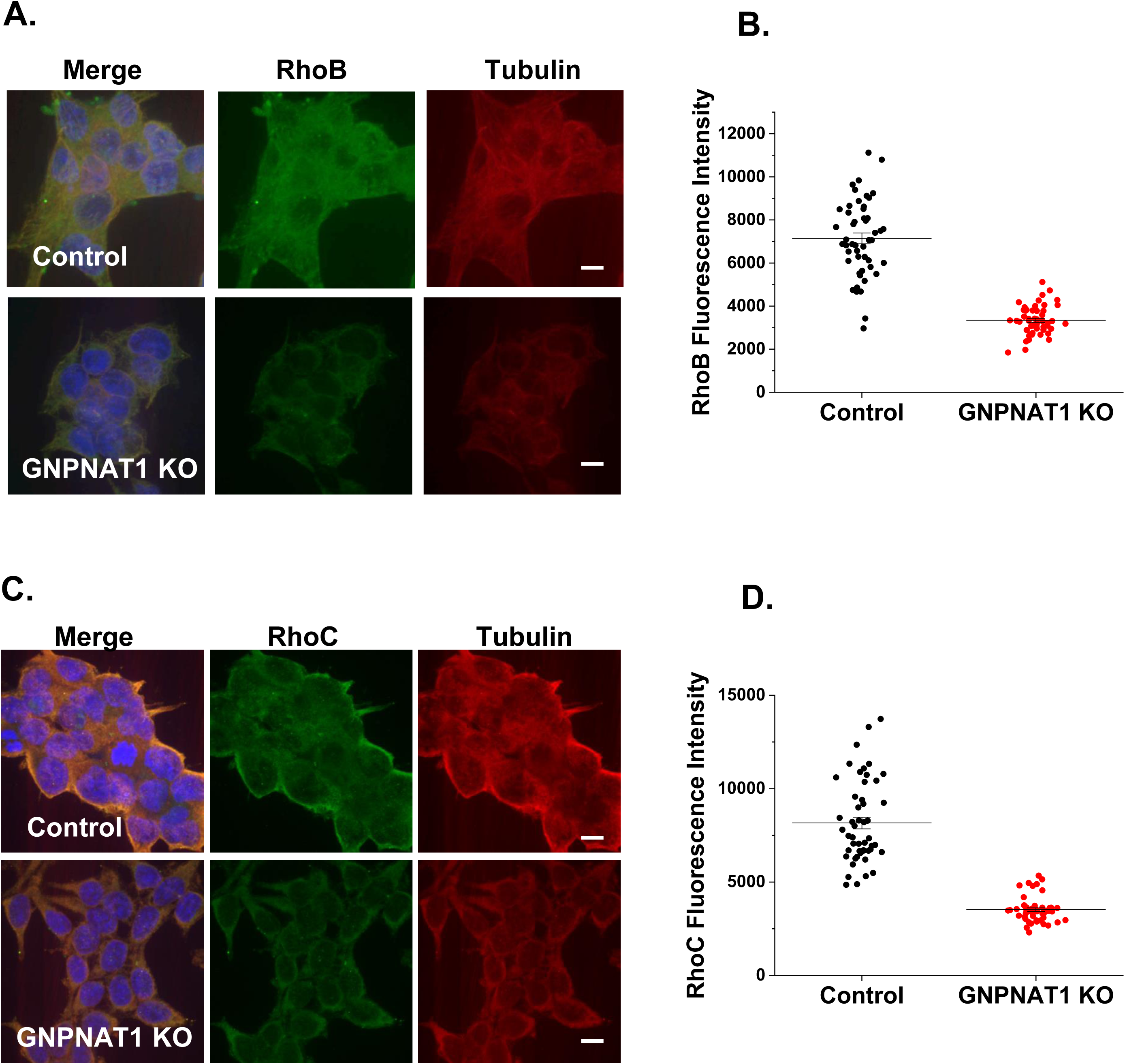
HBP depletion suppresses RhoB and RhoC levels. (A-D) Immunofluorescence staining (A and C) and the intensity quantification (B and D) of RhoB, and RhoC, respectively. Scale bars, 10 µm.

## References

1. Rawla P. Epidemiology of Prostate Cancer. World J Oncol [Internet]. 2019;10(2):63–89. Available from: http://www.wjon.org/index.php/WJON/article/view/1191

2. Harris WP, Mostaghel EA, Nelson PS, Montgomery B. Androgen deprivation therapy: Progress in understanding mechanisms of resistance and optimizing androgen depletion. Vol. 6, Nature Clinical Practice Urology. Nature Publishing Group; 2009. p. 76–85.

3. Chandrasekar T, Yang JC, Gao AC, Evans CP. Mechanisms of resistance in castration-resistant prostate cancer (CRPC). Transl Androl Urol. 2015;4(3):365–80.

4. Watson PA, Arora VK, Sawyers CL. Emerging mechanisms of resistance to androgen receptor inhibitors in prostate cancer. Vol. 15, Nature Reviews Cancer. Nature Publishing Group; 2015. p. 701–11.

5. DeWane G, Salvi AM, DeMali KA. Fueling the cytoskeleton-links between cell metabolism and actin remodeling. Vol. 134, Journal of Cell Science. Company of Biologists Ltd; 2021.

6. Liu G, Li J, Wu C. Reciprocal regulation of actin filaments and cellular metabolism. Eur J Cell Biol. 2022 Sep 1;101(4).

7. De Piano M, Manuelli V, Zadra G, Loda M, Muir G, Chandra A, et al. Exploring a role for fatty acid synthase in prostate cancer cell migration. Small GTPases. 2021;12(4):265–72.

8. Kaushik AK, Shojaie A, Panzitt K, Sonavane R, Venghatakrishnan H, Manikkam M, et al. Inhibition of the hexosamine biosynthetic pathway promotes castration-resistant prostate cancer. Nat Commun. 2016 May 19;7.

9. Parker AL, Kavallaris M, McCarroll JA. Microtubules and their role in cellular stress in cancer. Vol. 4 JUN, Frontiers in Oncology. Frontiers Research Foundation; 2014.

10. Akella NM, Ciraku L, Reginato MJ. Fueling the fire: Emerging role of the hexosamine biosynthetic pathway in cancer. Vol. 17, BMC Biology. BioMed Central Ltd.; 2019.

11. Paneque A, Fortus H, Zheng J, Werlen G, Jacinto E. The Hexosamine Biosynthesis Pathway: Regulation and Function. Vol. 14, Genes. Multidisciplinary Digital Publishing Institute (MDPI); 2023.

12. Muniz de Queiroz R, Oliveira IA, Piva B, Catão FB, Rodrigues B da C, Pascoal A da C, et al. Hexosamine Biosynthetic Pathway and Glycosylation Regulate Cell Migration in Melanoma Cells. Front Oncol. 2019;9(MAR).

13. Brooks Robey R, Weisz J, Kuemmerle N, Salzberg AC, Berg A, Brown DG, et al. Metabolic reprogramming and dysregulated metabolism: Cause, consequence and/or enabler of environmental carcinogenesis? Vol. 36, Carcinogenesis. Oxford University Press; 2015. p. S203–31.

14. Liu W, Jiang K, Wang J, Mei T, Zhao M, Huang D. Upregulation of GNPNAT1 Predicts Poor Prognosis and Correlates With Immune Infiltration in Lung Adenocarcinoma. Front Mol Biosci. 2021 Mar 25;8.

15. Yuan M, Breitkopf SB, Yang X, Asara JM. A positive/negative ion-switching, targeted mass spectrometry-based metabolomics platform for bodily fluids, cells, and fresh and fixed tissue. Nat Protoc. 2012;7(5):872–81.

16. Dimauro I, Pearson T, Caporossi D, Jackson MJ. A simple protocol for the subcellular fractionation of skeletal muscle cells and tissue. BMC Res Notes. 2012;5.

17. Goodson H V., Jonasson EM. Microtubules and microtubule-associated proteins. Cold Spring Harb Perspect Biol. 2018 Jun 1;10(6).

18. Dehmelt L, Smart FM, Ozer RS, Halpain S. The Role of Microtubule-Associated Protein 2c in the Reorganization of Microtubules and Lamellipodia during Neurite Initiation [Internet]. 2003. Available from: www.jneurosci.org

19. Liu Y, Ao X, Ding W, Ponnusamy M, Wu W, Hao X, et al. Critical role of FOXO3a in carcinogenesis. Vol. 17, Molecular Cancer. BioMed Central Ltd.; 2018.

20. Wang X, Hu S, Liu L. Phosphorylation and acetylation modifications of FOXO3a: Independently or synergistically? Vol. 13, Oncology Letters. Spandidos Publications; 2017. p. 2867–72.

21. Hanover G, Vizeacoumar FS, Banerjee SL, Nair R, Dahiya R, Osornio-Hernandez AI, et al. Integration of cancer-related genetic landscape of Eph receptors and ephrins with proteomics identifies a crosstalk between EPHB6 and EGFR. Cell Rep. 2023 Jul 25;42(7).

22. Pergaris A, Danas E, Goutas D, Sykaras AG, Soranidis A, Theocharis S. The clinical impact of the eph/ephrin system in cancer: Unwinding the thread. Vol. 22, International Journal of Molecular Sciences. MDPI AG; 2021.

23. Darling TK, Lamb TJ. Emerging roles for Eph receptors and ephrin ligands in immunity. Vol. 10, Frontiers in Immunology. Frontiers Media S.A.; 2019.

24. Pasquale EB. Eph receptors and ephrins in cancer: Bidirectional signalling and beyond. Vol. 10, Nature Reviews Cancer. 2010. p. 165–80.

25. Wang Q, Yang X, Xu Y, Shen Z, Cheng H, Cheng F, et al. RhoA/Rho-kinase triggers epithelial-mesenchymal transition in mesothelial cells and contributes to the pathogenesis of dialysis-related peritoneal fibrosis [Internet]. Vol. 9, Oncotarget. 2018. Available from: www.impactjournals.com/oncotarget/

26. Nishizuka M, Komada R, Imagawa M. Knockdown of RhoE Expression Enhances TGF-β-Induced EMT (epithelial-to-mesenchymal transition) in Cervical Cancer HeLa Cells. Int J Mol Sci. 2019 Oct 1;20(19).

27. Liu B, Dong H, Lin X, Yang X, Yue X, Yang J, et al. RND3 promotes Snail 1 protein degradation and inhibits glioblastoma cell migration and invasion [Internet]. Vol. 7, Oncotarget. 2016. Available from: www.impactjournals.com/oncotarget/

28. Bhowmick NA, Ghiassi M, Bakin A, Aakre M, Lundquist CA, Engel ME, et al. Transforming Growth Factor-1 Mediates Epithelial to Mesenchymal Transdifferentiation through a RhoA-dependent Mechanism. Vol. 12, Molecular Biology of the Cell. 2001.

29. Ren S, Shao Y, Zhao X, Hong CS, Wang F, Lu X, et al. Integration of metabolomics and transcriptomics reveals major metabolic pathways and potential biomarker involved in prostate cancer. Molecular and Cellular Proteomics. 2016 Jan 1;15(1):154–63.

30. Wellen KE, Lu C, Mancuso A, Lemons JMS, Ryczko M, Dennis JW, et al. The hexosamine biosynthetic pathway couples growth factor-induced glutamine uptake to glucose metabolism. Genes Dev. 2010 Dec 15;24(24):2784–99.

31. DeLiberty JM, Robb R, Gates CE, Bryant KL. Unraveling and targeting RAS-driven metabolic signaling for therapeutic gain. In: Advances in Cancer Research. Academic Press Inc.; 2022. p. 267–304.

32. Gómez-Cebrián N, Poveda JL, Pineda-Lucena A, Puchades-Carrasco L. Metabolic Phenotyping in Prostate Cancer Using Multi-Omics Approaches. Vol. 14, Cancers. MDPI; 2022.

33. Le Minh G, Esquea EM, Young RG, Huang J, Reginato MJ. On a sugar high: Role of O-GlcNAcylation in cancer. Vol. 299, Journal of Biological Chemistry. American Society for Biochemistry and Molecular Biology Inc.; 2023.

34. Logan CM, Menko AS. Microtubules: Evolving roles and critical cellular interactions. Vol. 244, Experimental Biology and Medicine. SAGE Publications Inc.; 2019. p. 1240–54.

35. Maccioni RB, V Cambiazo A. Role of Microtubule-Associated Proteins in the Control of Microtubule Assembly. Vol. 75, REVIEWS. 1995.

36. Bunning AR, Gupta ML. The importance of microtubule-dependent tension in accurate chromosome segregation. Vol. 11, Frontiers in Cell and Developmental Biology. Frontiers Media S.A.; 2023.

37. Vicente JJ, Wordeman L. The quantification and regulation of microtubule dynamics in the mitotic spindle. Vol. 60, Current Opinion in Cell Biology. Elsevier Ltd; 2019. p. 36–43.

38. Fong K wing, Leung JW chung, Li Y, Wang W, Feng L, Ma W, et al. MTR120/KIAA1383, a novel microtubule-associated protein, promotes microtubule stability and ensures cytokinesis. J Cell Sci. 2013 Feb;126(3):825–37.

39. Godek KM, Compton DA. Quantitative methods to measure aneuploidy and chromosomal instability. In: Methods in Cell Biology. Academic Press Inc.; 2018. p. 15–32.

40. Chen Y, Chen S, Li K, Zhang Y, Huang X, Li T, et al. Overdosage of Balanced Protein Complexes Reduces Proliferation Rate in Aneuploid Cells. Cell Syst. 2019 Aug 28;9(2):129–142.e5.

41. Holland AJ, Cleveland DW. Losing balance: The origin and impact of aneuploidy in cancer. Vol. 13, EMBO Reports. 2012. p. 501–14.

42. Sheltzer JM, Amon A. The aneuploidy paradox: Costs and benefits of an incorrect karyotype. Vol. 27, Trends in Genetics. 2011. p. 446–53.

43. Akhshi TK, Wernike D, Piekny A. Microtubules and actin crosstalk in cell migration and division. Vol. 71, Cytoskeleton. 2014. p. 1–23.

44. Coles CH, Bradke F. Coordinating Neuronal Actin-Microtubule Dynamics. Vol. 25, Current Biology. Cell Press; 2015. p. R677–91.

45. Tang DD, Gerlach BD. The roles and regulation of the actin cytoskeleton, intermediate filaments and microtubules in smooth muscle cell migration. Vol. 18, Respiratory Research. BioMed Central Ltd.; 2017.

46. Zhang X, Tang N, Hadden TJ, Rishi AK. Akt, FoxO and regulation of apoptosis. Vol. 1813, Biochimica et Biophysica Acta - Molecular Cell Research. 2011. p. 1978–86.

47. Dan HC, Antonia RJ, Baldwin AS. PI3K/Akt promotes feedforward mTORC2 activation through IKKα [Internet]. Vol. 7. Available from: www.impactjournals.com/oncotarget

48. Dan HC, Ebbs A, Pasparakis M, Van Dyke T, Basseres DS, Baldwin AS. Akt-dependent activation of mTORC1 complex involves phosphorylation of mTOR (mammalian target of rapamycin) by IκB kinase α (IKKα). Journal of Biological Chemistry. 2014;289(36):25227– 40.

49. Fu W, Hall MN. Regulation of MTORC2 signaling. Vol. 11, Genes. MDPI AG; 2020. p. 1–19.

50. Saxton RA, Sabatini DM. mTOR Signaling in Growth, Metabolism, and Disease. Vol. 168, Cell. Cell Press; 2017. p. 960–76.

51. Black AR, Black JD. Protein kinase C signaling and cell cycle regulation. Vol. 3, Frontiers in Immunology. 2012.

52. Hsu AH, Lum MA, Shim KS, Frederick PJ, Morrison CD, Chen B, et al. Crosstalk between PKCα and PI3K/AKT Signaling Is Tumor Suppressive in the Endometrium. Cell Rep. 2018 Jul 17;24(3):655–69.

53. Liu M, Clarke CJ, Salama MF, Choi YJ, Obeid LM, Hannun YA. Co-ordinated activation of classical and novel PKC isoforms is required for PMA-induced mTORC1 activation. PLoS One. 2017 Sep 1;12(9).

54. Saxton RA, Sabatini DM. mTOR Signaling in Growth, Metabolism, and Disease. Vol. 168, Cell. Cell Press; 2017. p. 960–76.

55. Kalluri R, Weinberg RA. The basics of epithelial-mesenchymal transition. Vol. 119, Journal of Clinical Investigation. 2009. p. 1420–8.

56. Chen T, You Y, Jiang H, Wang ZZ. Epithelial–mesenchymal transition (EMT): A biological process in the development, stem cell differentiation, and tumorigenesis. Vol. 232, Journal of Cellular Physiology. Wiley-Liss Inc.; 2017. p. 3261–72.

